# The Major-Minor mode Dichotomy in Music Perception: A Systematic Review and Meta-Analysis on its Behavioural, Physiological, and Clinical Correlates

**DOI:** 10.1101/2023.03.16.532764

**Authors:** Giulio Carraturo, Victor Pando-Naude, Marco Costa, Peter Vuust, Leonardo Bonetti, Elvira Brattico

## Abstract

Since ancient Greece, major and minor modes in Western tonal music have been identified as the primary responsible musical feature for eliciting emotional states. As such, the underlying correlates of the major-minor mode dichotomy in music perception have been extensively investigated through decades of psychological and neuroscientific research, providing plentiful yet often discordant results. Specifically, crucial questions remain about the several factors contributing to the affective perception of major and minor modes, at times very different among individuals. Moreover, major and minor mode perception has never been quantitatively compared in literature. This comprehensive systematic review and meta-analysis aimed to provide a qualitative and quantitative synthesis of musical mode perception and its behavioural and neural correlates. The qualitative synthesis resulted in 69 studies, showing great diversity in how the major-minor dichotomy has been empirically approached. Most studies reviewed were conducted on adults, considered participants’ expertise, employed real-life musical stimuli, performed behavioural evaluations, and were carried out among Western listeners. Behavioural, electroencephalography, and neuroimaging meta-analyses (36 studies) consistently showed that major and minor mode elicit distinct neural and emotional responses. Based on our findings, a framework to describe a *Major-Minor Mode(l)* of music perception and its behavioural and physiological correlates is proposed, incorporating individual factors such as age, expertise, cultural background, and emotional disorders. Limitations, implications, and suggestions for future research are discussed, including putative clinical applications of major-minor dichotomy and best practices regarding stimulation paradigms for musical mode investigation.

**Public Significance Statement:** This study provides qualitative and quantitative evidence of the distinct behavioral and neural responses elicited by major and minor mode, while also highlighting the influence of factors such as age, culture, personality, and health. Results offers a detailed overview of the major-minor dichotomy in music, putting forward an integrated and critical discussion of methodologies, paradigms, and clinical implications of this pivotal musical feature.

---

A fundamental dichotomy in Western tonal music, linked to the expression of emotional valence in music, is that between major and minor modes. Musical mode is determined by the pattern of tones and semitones in its scales, and the difference between major and minor modes is rooted in the interval between the tonic note and the third degree of the scale. For instance, in C major, the third degree is an E, whereas in C minor the third degree is one semitone lower, i.e., an E flat. Minor mode is expressed with three different scales: natural minor, melodic minor, and harmonic minor. In addition, the modern Dorian mode is often used as a minor mode (**Figure 1**).

**Figure 1.**
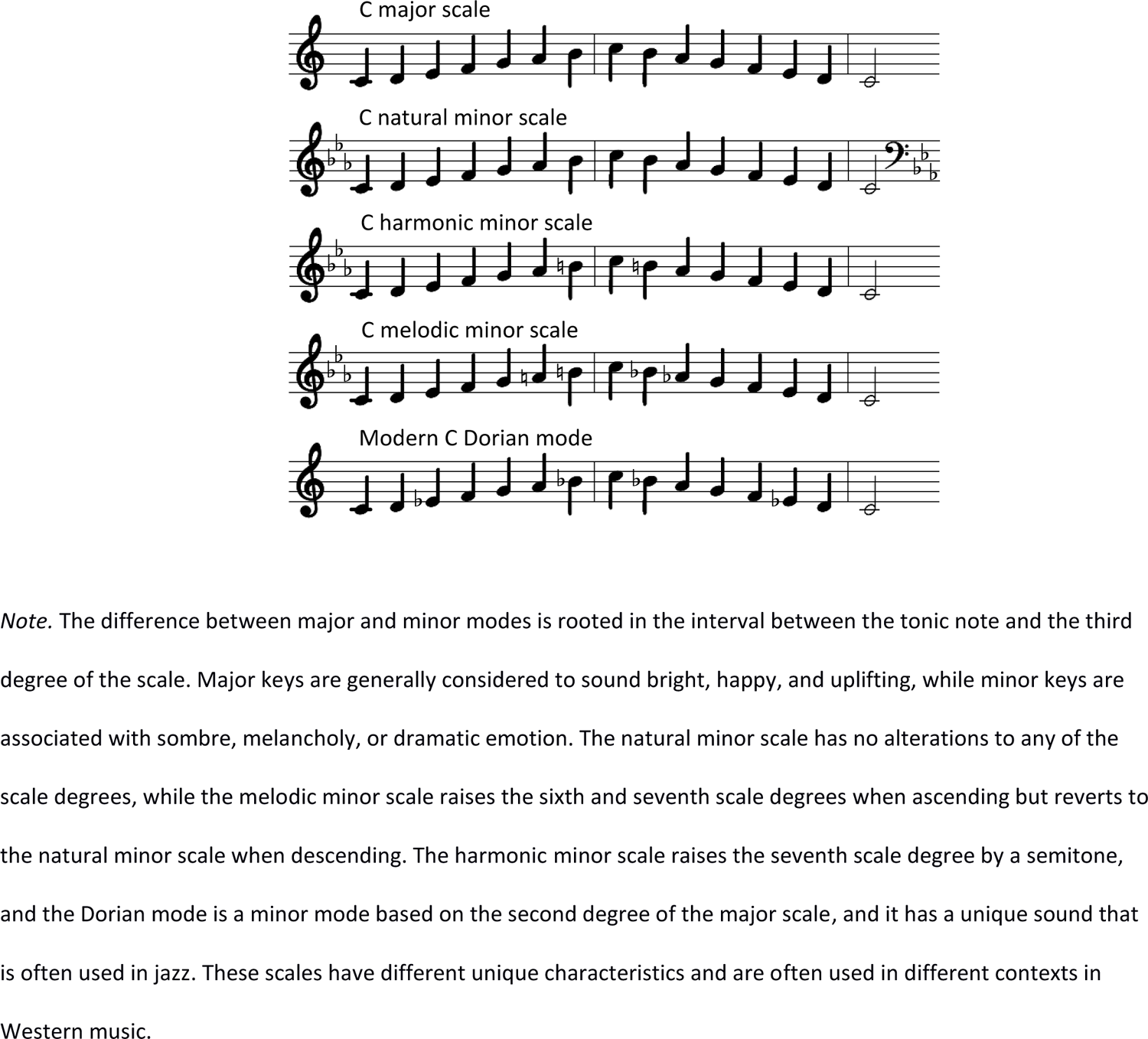
The Major and Minor Modes in Western Tonal Music

Major keys are generally considered to sound bright, happy, and uplifting, while minor keys are associated with sombre, melancholy, or dramatic emotion. However, these associations are not always straightforward or universal, and composers have used both major and minor modes to convey a wide range of emotions and affective states (Chubb et al., 2013; Virtala & Tervaniemi, 2017).

Regarding the different types of minor modes in Western tonal music, each has its own unique characteristics and are often used in different contexts. For example, the natural minor scale is the most basic form of minor scale, and it is simply a minor scale with no alterations to any of the scale degrees. The melodic minor scale, on the other hand, raises the sixth and seventh scale degrees when ascending, but reverts to the natural minor scale when descending. This creates a slightly different sound and allows for more flexibility in melodic writing. The harmonic minor scale, meanwhile, raises the seventh scale degree by a semitone, which creates a strong pullback to the tonic note and is often used in harmonic progression. Finally, the Dorian mode is a minor mode that is based on the second degree of the major scale, and it has a unique sound that is often used in jazz and other styles of music (Casas-Mas et al., 2015).

Western listeners are generally familiar with the major-minor tonal system due to its prevalence in everyday music (Koelsch, 2009). Major and minor are indeed the most used modes in Western music, and their structural difference produces prominent distinct emotional responses. Therefore, mode is considered, together with tempo, a strong determinant of musical-emotional responses (Peretz et al., 1998). Some authors argued that major-minor dichotomy is actually the main cue contributing to emotional expression in music (Eerola et al., 2013). Notably, the explicit emotional recognition of music mode develops later in life than the recognition of tempo, and appears to depend on both individual characteristics and cultural factors, especially in the first years of life (Bonetti & Costa, 2019; Dalla Bella et al., 2001). Previous studies have suggested that musical training and age can further modify major-minor perception and its affective processing (Castro & Lima, 2014; Virtala & Tervaniemi, 2017), even though no explicit knowledge of major-minor mode structures is needed in order to produce the related emotional responses (Pallesen et al., 2003). However, there is evidence of large and systematic inter-individual differences, since several factors such as expertise, mood, and personality may affect major-minor perception and preference (Bonetti & Costa, 2016; Brattico et al., 2016; Chubb et al., 2013; Hunter et al., 2011; Ladinig & Schellenberg, 2012).

Neuroscientific studies, such as electroencephalography (EEG), magnetoencephalography (MEG), positron emission tomography (PET), and functional magnetic resonance imaging (fMRI), have revealed that major and minor modes of music activate distinct brain regions, with a significantly stronger involvement of limbic structures and reward circuits in response to minor mode (Green et al., 2008; Suzuki et al., 2008). However, the findings of these studies are heterogeneous, likely due to differences in methodology and type of stimulation. Interestingly, specific neural activity in response to music mode can be detected already in the very first years of life, as indexed by EEG measures studies (Virtala et al., 2013), whereas divergent results have been observed among older children without music training (Virtala et al., 2012). Additionally, several studies report biased emotional processing of musical features (including musical mode) in both clinical and subclinical conditions (Bonetti et al., 2017; Gosselin, 2005; Punkanen et al., 2011; Zhao et al., 2019).

There is a vast literature on the behavioural, brain, and physiological correlates of the major-minor dichotomy in music perception. However, the variety of methodological approaches used has hindered a unifying view and has led to contradictory results. Moreover, previous reviews have explored the behavioural and physiological correlates of other musical features such as rhythm, tempo, pitch, timbre, melody, harmony, and syntax (Koelsch, 2011; Peretz & Zatorre, 2005). In terms of musical mode, Virtala & Tervaniemi (2017) summarised evidence on the perception of musical mode focusing mainly on the neurocognitive basis of this dichotomy, and suggested that major-minor and consonance/dissonance processing are present early in development and have partly universal, biologically hardwired origins, likely related to human vocalisations (Cook et al., 2006). Also, brain maturation, musical enculturation, and music training significantly modulate sensory and affective processing. More recently, Asano et al. (2022) conducted a meta-analysis of neuroimaging studies focusing on tonal cognition, with an emphasis on harmony processing, and identified specific brain areas in the right frontal lobe that are similar to those involved in affective speech prosody.

Despite the extensive literature exploring major-minor dichotomy and its centrality in Western music, no study has thus far provided a systematic and unified investigation of this feature or on how its perception is affected by person-related factors. To this end, we conducted a systematic review to comprehensively summarise and organise the literature on major-minor mode perception and its behavioural and brain correlates, by performing both a qualitative and quantitative synthesis. Specifically, we aimed to answer the following questions: what factors influence individual sensitivity and affective perception of major-minor dichotomy? What types of methods and paradigms are used to investigate it? What are the brain mechanisms and neural structures underlying major-mode perception? Can musical mode be seen as a clinical stimulation paradigm? What are the implications of the findings for future research? The findings are expected to significantly contribute to the psychology of music advancing knowledge on a major determinant of affective and aesthetic responses to music.

## Methods

### Search Strategy

This systematic review followed procedures form the Cochrane Handbook for Systematic Reviews (Higgins & Green, 2011) and from the Centre for Reviews and Dissemination (Culyer, 2014). The systematic approach was carried in accordance with the PRISMA statement (Moher et al., 2009). Three authors (GC, VPN, and MC) each performed a search in the electronic databases Scopus, PubMed and PsycInfo by combining the following keywords and MeSH terms opportunely adapted to each database: music, musician, musical mode, music perception, music performance, music playing, music composition, major mode, minor mode, musical training, expertise, behavioural correlates, individual differences, cognition, emotion, happiness, sadness, valence, arousal, mood disorders, depression, neural correlates, auditory stimulation, magnetic resonance imaging, magnetoencephalography, electroencephalography, evoked potentials, transcranial magnetic stimulation, electrocorticography (details in **Supplementary Information**). The reference lists of the included articles were scrutinised manually for additional publications. The search results were imported to *Covidence*, in RIS or XML format, for screening.

### Study Selection

Three authors (GC, VPN, and MC) independently and systematically screened the literature by titles and abstracts and then read the full texts to identify their eligibility using the Covidence tool. For the qualitative synthesis, studies investigating musical mode perception, including all ages and levels of musical expertise, in English, and published in refereed journals were selected. Also, studies in which musical mode was not the main aim of investigation but that offered relevant results to our purpose were included, as well as studies exploring musical mode perception considering clinical conditions. Studies not in English, not including stimuli in both major and minor mode, single cases, reviews, and dissertations were excluded. No years or places of publication were imposed. For the quantitative synthesis (meta-analysis), studies were included if reported any measurement (i.e., behavioural or physiological) comparing positive (happy, pleasant, like) and negative (sad, unpleasant, dislike) emotional connotations. Any disagreements in the literature search and data extraction were resolved by a fourth author (LB). As for the methods used in the reviewed studies, “physiological” refers to methods of brain imaging and brain neurophysiology, as well as psychophysiological measurements such as heart rate and skin conductance; “behavioural” refers to tasks usually involving the affective evaluation of musical stimuli.

### Data Extraction

For the qualitative synthesis, data extraction was performed by one author (GC) and cross-checked by other two authors (VPN and MC). The following information was extracted from each study: year of publication, geographical location, number of participants, age group (adults and/or children), musical expertise (musicians, non-musicians, undefined), type of stimulation (chords, excerpts, musical pieces, and/or tone scrambles), centrality of musical mode in the research questions of each study, and type of method (i.e., behavioural, EEG, MEG, PET). The great variety of stimuli used among the studies was extracted and labelled for each study as “chords” when the stimulation was a single or a short sound (maximum 2s duration), “excerpts” with musical stimuli ranging from 2.5s to 29.5s, or “musical pieces” for all those of 30s or more. Finally, the fourth category of stimuli consists of “tone scrambles”: rapid, randomly ordered sequences of pure tones. For the quantitative synthesis, effect sizes and standard errors were extracted from behavioural and EEG studies, and stereotactic coordinates were extracted from fMRI and PET studies in either Talairach or Montreal Neurological Institute (MNI) three-dimensional-coordinate system.

### Quality Assessment

As there is no universally accepted and evidence-based tool available to evaluate the quality of studies across different research designs, the task of quality assessment becomes challenging. To comprehensively assess the quality of the included studies in this systematic review, the QualSyst tool (Kmet et al., 2004) was used. This tool offers a comprehensive and quantitative approach to evaluate the quality of research, covering a diverse range of study designs, in a systematic and reproducible manner. A summary score is calculated for each paper by summing the total score obtained across relevant items and dividing by the total possible score (i.e.: 28 – (number of “n/a” x 2)). A score higher than 0.75 suggests high quality, between 0.55 and 0.75 suggests moderate quality, and lower than 0.55 suggests low quality in the study.

### Meta-analyses

Meta-analyses were carried out for behavioural studies, EEG, and fMRI studies, independently. For behavioural and EEG studies, meta-analyses of effect sizes were performed (Harrer et al., 2021). For fMRI studies, coordinated-based meta-analyses (CBMA) were conducted using the activation likelihood activation (ALE) method (Eickhoff et al., 2009). The studies covered various paradigms, including “Like-dislike,” “Pleasant/unpleasant,” “Happiness/sadness,” with positive emotions in major mode and negative emotions in minor mode, and assessed emotions in terms of whether they were “Felt” or “Perceived.”

### Behavioural and EEG Meta-analyses

For behavioural and EEG studies, effect sizes were used to conduct meta-analyses. Additionally, standard errors were extracted from studies to account the standard deviation of the sampling distribution. To interpret the magnitude of the observed effects, effect sizes of 0.2 were considered small, 0.5 were considered medium, and 0.8 were considered large. To order, calculate, and compare effect sizes, this study followed the recommendations by Harrer et al., 2021. The effect sizes corresponded to the comparison between positive (happy, pleasant, like) and negative (sad, unpleasant, dislike) emotional connotations. Effect sizes and/or standard errors were calculated from other statistical analyses (e.g., mean and standard deviation, correlation, ANOVA, etc.) if a given study did not report it specifically.

Random-effects models were employed since variation in effect sizes among studies was assumed to occur because of random sampling error as well as differences between groups or individuals (Cooper & Hedges, 1994). This approach allowed for generalizations about the effects across a population rather than just past studies (Raudenbush, 1994). To address small-sample bias, a correction was applied by weighting the effect size associated with studies by sample size (Hedges & Olkin, 1985). The random-effects meta-analysis was performed using the metagen function with the Hartung-Knapp (HK) adjustment for the random-effects model, considering the potential heterogeneity among studies. Additionally, the Sidik-Jonkman (SJ) estimator was used for tau^2 and tau, and the Q-Profile method was used to compute confidence intervals for these estimates. Heterogeneity among the included studies was quantified using the I^2 statistic, which measures the proportion of total variation attributable to heterogeneity rather than sampling error. The Q statistic test was applied to assess the significance of heterogeneity. A p-value less than 0.05 was considered indicative of significant heterogeneity.

To estimate the uncertainty of the meta-analysis results, prediction intervals were calculated based on t-distribution. Prediction intervals provide an estimate of the range within which the true effect size of a future study is likely to fall. The results were presented using descriptive statistics, including the pooled effect size (SMD) and its 95% Confidence Interval (CI). The Z-value and p-value were reported to assess the significance of the overall effect. Additionally, the tau^2 and tau estimates were provided as measures of between-study variance and standard deviation of study effects, respectively. Forest plots were generated to visualize the individual study effect sizes along with their confidence intervals and the pooled effect size estimate. A subgroup analysis in the behavioural meta-analysis was conducted to test the effect of felt vs perceived emotion.

### fMRI and PET studies

In this study, the GingerALE software v3.0.2 (Eickhoff et al., 2009; Fox et al., 2013, 2014) was utilized to conduct CBMAs using the ALE method from the BrainMap platform (Fox et al., 2005). Coordinates (foci) from each study were extracted, and if required, converted from Talairach to MNI space using the Lancaster transform (icbm2tal) provided in GingerALE (Eickhoff et al., 2012; Laird et al., 2010). The ALE method assesses anatomical consistency and concordance among studies by weighting coordinates based on sample size, producing estimates of anatomical likelihood for each intracerebral voxel on a standardized brain map.

The algorithm treats anatomical foci as spatial probability distributions centred at the specified coordinates, evaluating the correlation between the spatial locations of foci across MRI studies investigating the same construct, and comparing it to a null distribution of random spatial association between experiments. Statistical significance of the ALE scores was determined using a permutation test with cluster-level inference at a threshold of p < 0.05 (FWE), and a cluster-forming threshold set at p < 0.001. Separate ALE meta-analyses were conducted for results of major > minor and minor > major, independently.

The BrainMap platform is a large-scale database that stores findings from neuroimaging studies that can be used with a meta-analytic approach by exploring the co-activation of brain regions with a specific region-of-interest (ROI) across different tasks, rather than on a particular one. Meta-analytic connectivity modelling (MACM) identifies the functional network of the ROI, i.e., its functional connectivity (FC). MACM allows for the functional segregation of each ROI’s potential contribution to different behavioural domains (Laird et al., 2013; Robinson et al., 2010). To conduct the co-activation analyses, Sleuth (Laird et al., 2009) and GingerALE, both part of the BrainMap platform, were used. For the identification of significant convergence areas, an ALE meta-analysis was performed on all foci obtained after individually searching Sleuth for each ROI. The search criteria included “context: normal mapping” and “activations: activation only”. ROIs were created using Mango with a 5mm-radius sphere on the MNI152_T1_1mm brain template. The results of each ROI search were then exported to GingerALE, where a permutation test was conducted using cluster-level inference at p < 0.05 (FWE), with a cluster-forming threshold set at p < 0.001 to identify regions of significance. To further understand the functional characteristics of the ROIs, the study utilized the “Behavioural Domain” meta-data categories available in the BrainMap platform, including action, perception, emotion, and interoception.

CBMA’s, such as ALE, are valuable tools for integrating findings from multiple studies to draw meaningful conclusions about brain activation patterns. However, they are not immune to certain biases inherent in all meta-analyses, such as publication bias, which could distort results and undermine the validity of the conclusions drawn. One significant concern is the “file drawer problem,” where studies with non-significant or null results are less likely to be published, leading to an overrepresentation of studies with positive findings. To address potential publication bias and assess the robustness of the results, the Fail-Safe N analysis (FSN) was performed (Acar et al., 2018). FSN evaluates how much opposing evidence would need to be added to a meta-analysis before the findings change significantly. For instance, a typical 95% confidence interval for the number of studies reporting no local maxima (effectively null findings) in normal human brain mapping studies varies from 5 to 30 per 100 published studies. Taking the upper bound (30%) and considering that the ALE meta-analyses in this study included 7 experiments in the major-minor and 8 experiments in the minor-major ALE meta-analyses, the minimum FSN was set at 2.

### Transparency and Openness

We adhered to the PRISMA 2020 guidelines for systematic reviews (Page et al., 2021). All data and research materials (including our coding scheme) are available at the Supplemental Material. The data supporting the findings of this study is freely available at the Open Science Framework (OSF) website: https://osf.io/76kmb/?view_only=c59b316f6ce646f49b9f8440e5a98056

## Results

### Characteristics of Included Studies

The PRISMA flowchart for systematic reviews (**Figure S1**) and the PRISMA checklist (**Table S1**) provide a thorough overview of the identification, screening, eligibility, and inclusion procedure of studies. A total of 2,406 records were identified through manual and database searching. After removing duplicates, 1,858 articles were initially screened by title and abstract. Next, 386 records were screened by full-text review. Finally, 69 publications were included in the qualitative synthesis, with a total of 4,441 subjects.

In **Table 1** we listed all the studies included in our systematic review and their characteristics. The geographical distribution of the studies is: USA (12), Canada (9), Finland (9), Japan (8), Denmark (6), UK (6), Italy (5), China (2), Australia (2), Russia (2), France (2), Netherlands (1), Portugal (1), Croatia (1), Switzerland (1), Taiwan (1), and Germany (1). This information highlights how studies investigating musical mode perception have been carried out mainly among Western listeners. All in all, most studies (63%) have been published from 2010 onwards. Thirty-eight articles specifically aimed at major-minor mode investigation, whereas in the remaining 31 studies this dichotomy is included but is not the specific focus.

**Table 1.**
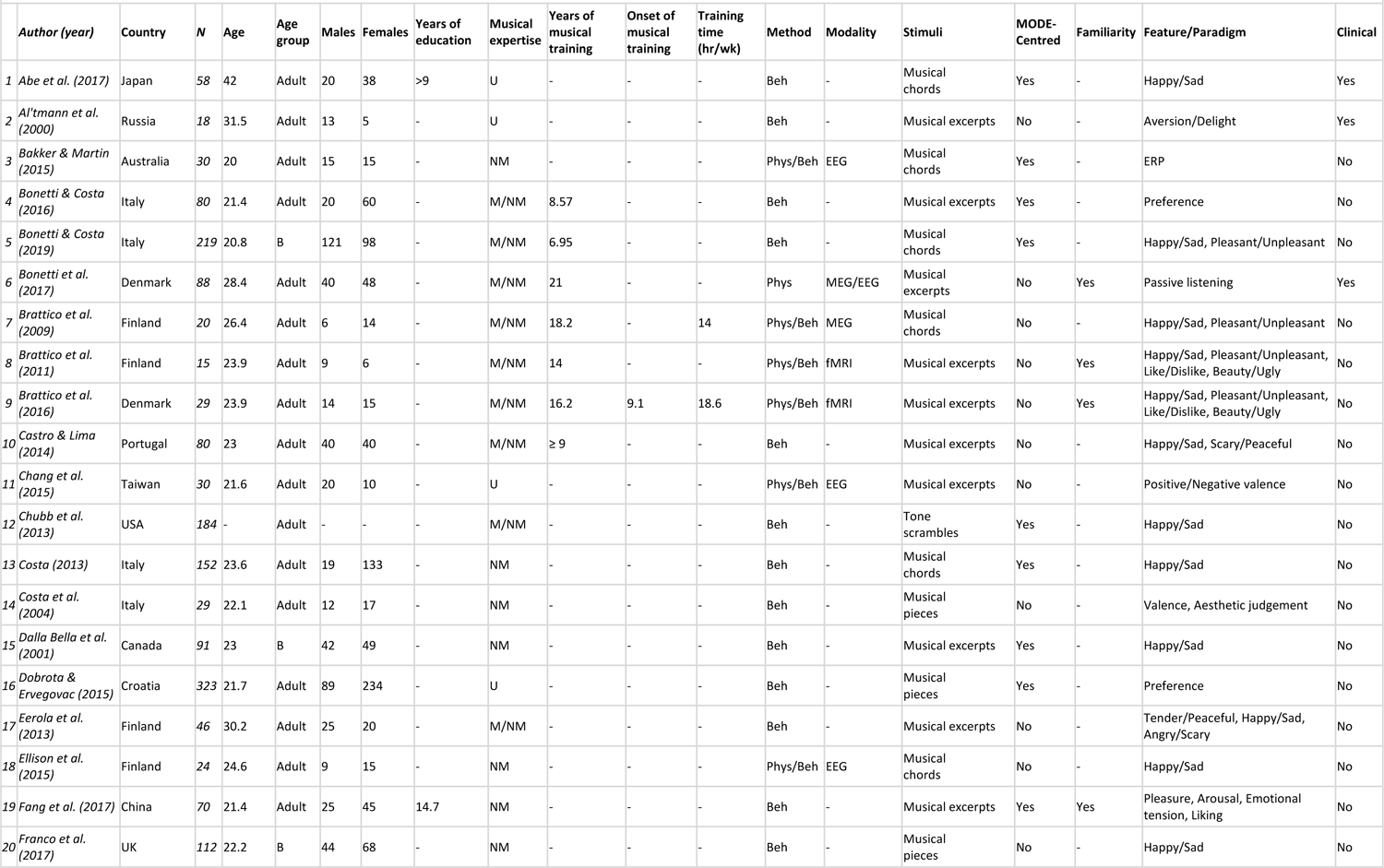
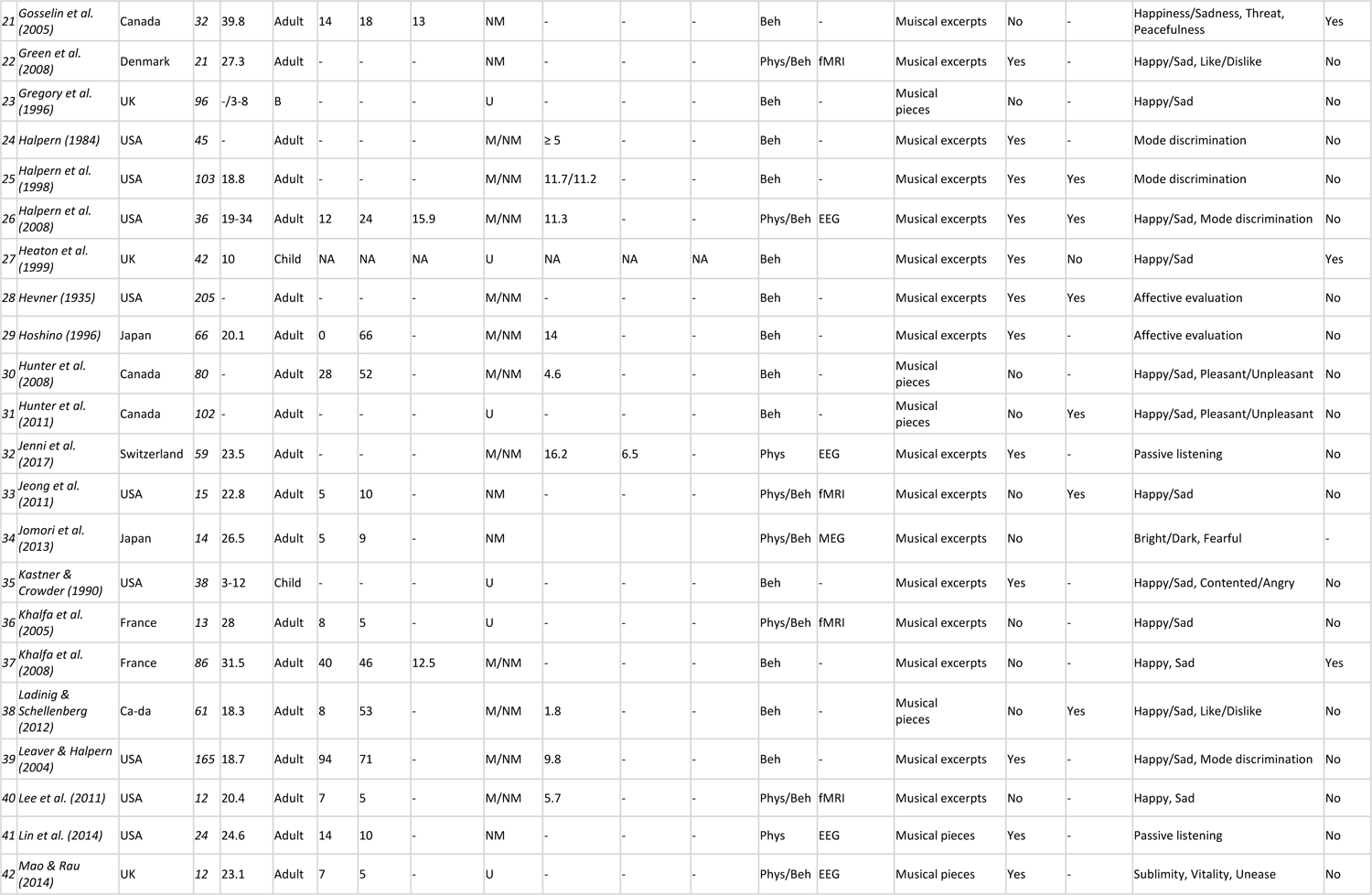
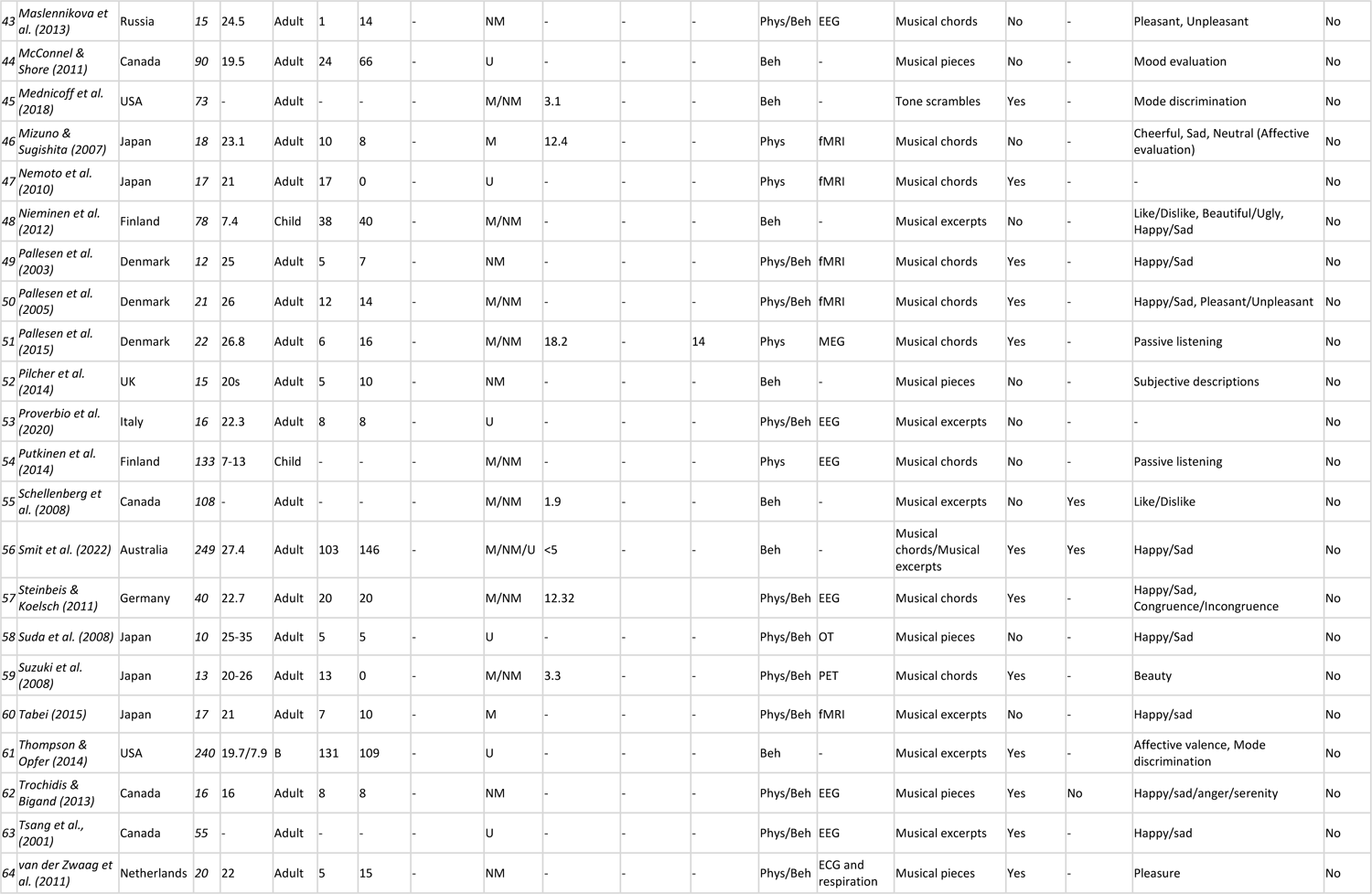
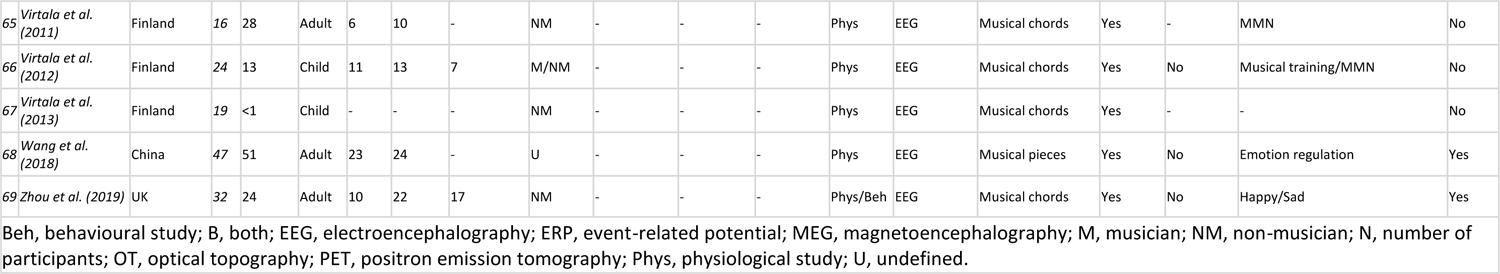
Characteristics of included studies.

As visible from **Table 2**, which contains a cross-tabulation of the studies as a function of the method and the sample, we obtained a larger body of empirical research investigating the major-minor dichotomy in the adult population (58) as compared to research conducted with children (6), with only 5 studies including both groups. Most of the studies (52 out of 69) reported participants’ level of musical expertise, although the years of musical training were indicated only in 24 articles (including 4 papers also reporting training time per day/week). Participants’ level of education in years was described in very few articles (7 out of 69). Behavioural studies represent most articles reviewed here (33), whereas a few applied exclusively physiological measures (10). The remaining 26 papers applied both physiological and behavioural methods. Studies considering clinical samples are 8, including one study on subclinical population. Finally, we summarised all the main findings relevant to the questions addressed in **Table S2**.

**Table 2.** Cross-tabulation of the Studies as a Function of Method, Expertise, and Sample.

|  | Physiological | Behavioural | Physiological &<br>Behavioural |
| --- | --- | --- | --- |
| Children | 3 | 3 | 0 |
| Adults | 8 | 25 | 25 |
| Children & Adults | 0 | 5 | 0 |
| Musicians | 1 | 0 | 1 |
| Non-musicians | 3 | 7 | 10 |
| Musicians & Non- musicians | 5 | 17 | 8 |
| Undefined expertise | 2 | 9 | 6 |

### Types of Stimulation

In **Table 1**, we indicated the type of stimulation as “chords”, “excerpts”, “musical pieces” or “tone scrambles”. This distinction was necessary considering the variety of stimuli and their durations used in the studies. For instance, some of the studies reviewed employed musical chords (usually less than 1s) (Pallesen et al., 2015; Putkinen et al., 2014; Steinbeis & Koelsch, 2011; Virtala et al., 2012), whereas among other investigations, the eliciting stimuli were longer than 5 minutes (McConnell & Shore, 2011; van der Zwaag et al., 2011), or composed of “non-musical” tone sequences (Chubb et al., 2013; Mednicoff et al., 2018). Such methodological differences needed to be identified and considered in the evaluation of the elicited responses. Included in this review are 18 studies using musical chords as stimuli, 32 studies employing musical excerpts, 15 with longer-lasting musical stimuli/musical pieces, 2 including both chords and excerpts, and 2 employing tone scrambles. Arguably, the behavioural and physiological responses elicited by chords or short fragments of music are more likely to be a direct consequence of major-minor dichotomy. Longer real-life music pieces in turn are more ecologically valid since they are more similar to the type of music to which we are exposed every day. However, in the latter, mode may be confounded with other aspects of musical structure conveying emotions.

### Centrality of Musical Mode

Most of the papers reviewed consist of investigations specifically aimed at major-minor mode (54%). However, we decided to also include studies not aimed exclusively at investigating this dichotomy since it is quite common that more than one feature is simultaneously investigated within the same study (such as mode, timbre, and tempo); yet we considered the role of our keyword (musical mode) significant for our questions. The “indirect” role of the keyword could be explained considering that several studies mainly focused on investigating emotional responses to music rather than major-minor dichotomy per se. In this perspective, the distinct affective connotation associated with major-minor mode was often exploited as an inducer of happy/sad emotional responses. Thus, in these studies musical mode was the independent variable, and the emotional response was the dependent variable.

### Quality Assessment

The quality of the included studies in this review was comprehensively assessed with the QualSyst tool following the recommendations by Kmet et al. (2004). Since we did not include any qualitative studies in this review, only the checklist for assessing the quality of quantitative studies was used. Additionally, the review only included observational studies, and no interventional studies, and therefore items 5, 6, and 7 were scored as not applicable (NA). The total scores for each study ranged from 0.32 to 1.0 with a mean of 0.9 ± 0.12. Overall, the included studies showed good practices with high scores in items related to design description and interpretation of results and conclusions; lower scores in items related to subject characteristics and controlling for confounding effects (**Table S3**). To ensure adequate quality in the neuroimaging studies included in meta-analyses, **Table S4** shows fMRI acquisition and analysis information.

### Meta-analysis of Behavioural Studies

The behavioural meta-analysis conducted using the random effects model encompassed a total of 12 experiments, after excluding one outlier (Green et al., 2008), investigating the relationship between musical mode and emotional response. The meta-analysis revealed a significant pooled random effects SMD = 0.2167 (95%-CI [0.1089, 0.3245], *p* = 0.001), and a prediction interval for the effect size ranged from −0.2341 to 0.6675. Quantifying heterogeneity, the analysis found a tau^2 of 0.0360 (95%-CI [0.0011 to 0.0696]), indicating substantial variability among the included studies. The I^2 statistic indicated moderate-to-high heterogeneity (I^2 = 65.3%). The test of heterogeneity (Q-test) was statistically significant (Q = 31.66, df = 11, p < 0.001), further supporting the presence of heterogeneity among the studies. The leave-one-out analysis demonstrated that the effect size remained robust upon excluding individual studies. A forest plot was generated to visually represent the effect sizes and confidence intervals for each study (**Figure 2**). The subgroup analysis comparing felt vs perceived emotion did not yield significant results.

**Figure 2.**
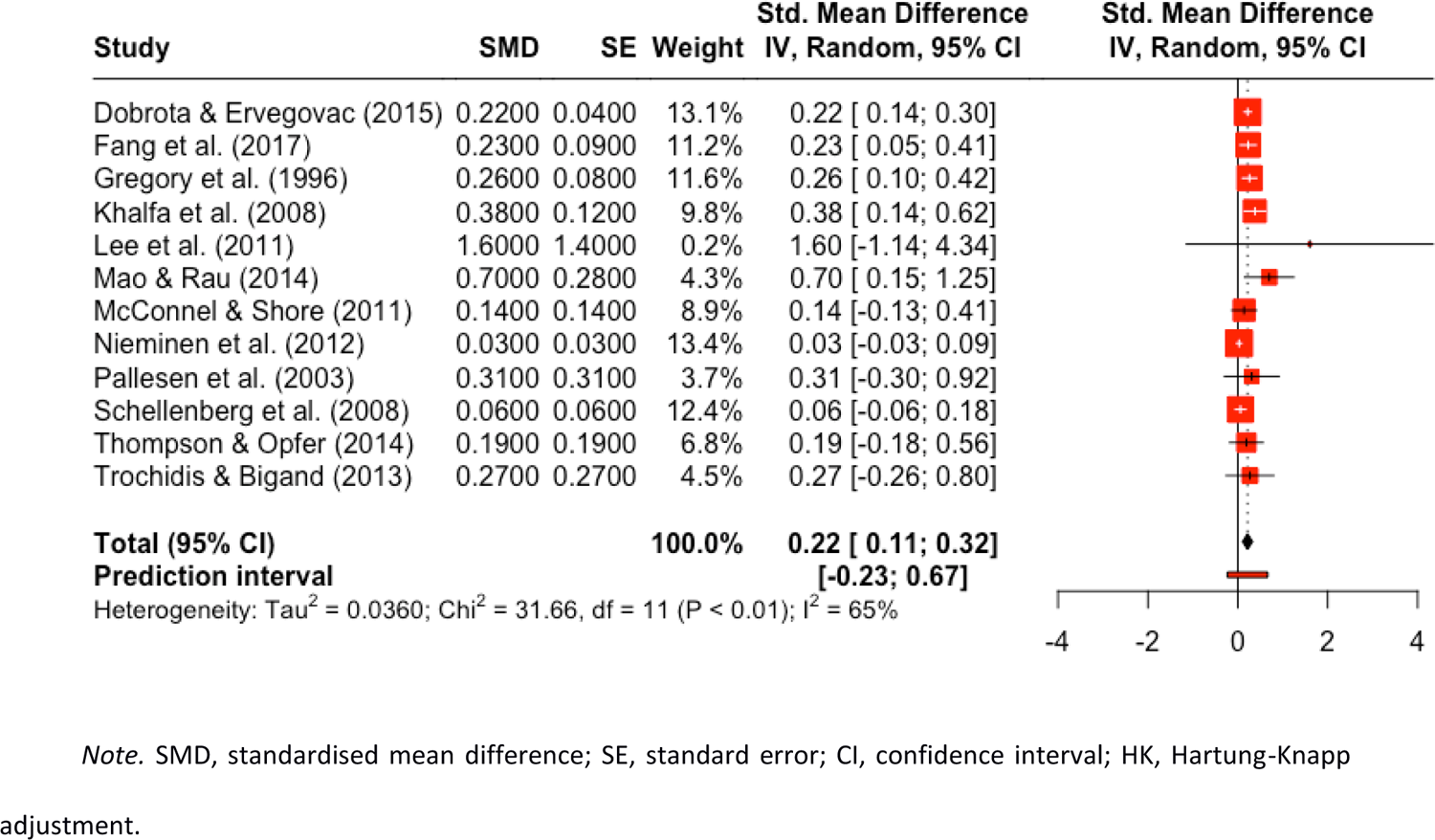
Behavioral Meta-analysis forest plot.

### Meta-analysis of Electroencephalography Studies

The EEG meta-analysis conducted using the random effects model encompassed a total of 12 experiments. A significant pooled SMD = 0.1621 (95% CI [0.0913, 0.2328], *p* = 0.0004) was found, indicating a moderate effect of musical mode on neural activity, presumably driven by emotional connotations. The prediction interval for the SMD ranged from −0.1098 to 0.4339, signifying the potential effect sizes that may be observed in future similar studies. Heterogeneity among the studies was low (I² = 0.0%), suggesting consistent findings across the diverse dataset. A test of heterogeneity (Q = 6.35, df = 11, p = 0.8493) showed no statistically significant differences in effect sizes among the studies. The leave-one-out analysis demonstrated that the effect size remained robust upon excluding individual studies. No outliers were detected. A forest plot was generated to visually represent the effect sizes and confidence intervals for each study (**Figure 3**).

**Figure 3.**
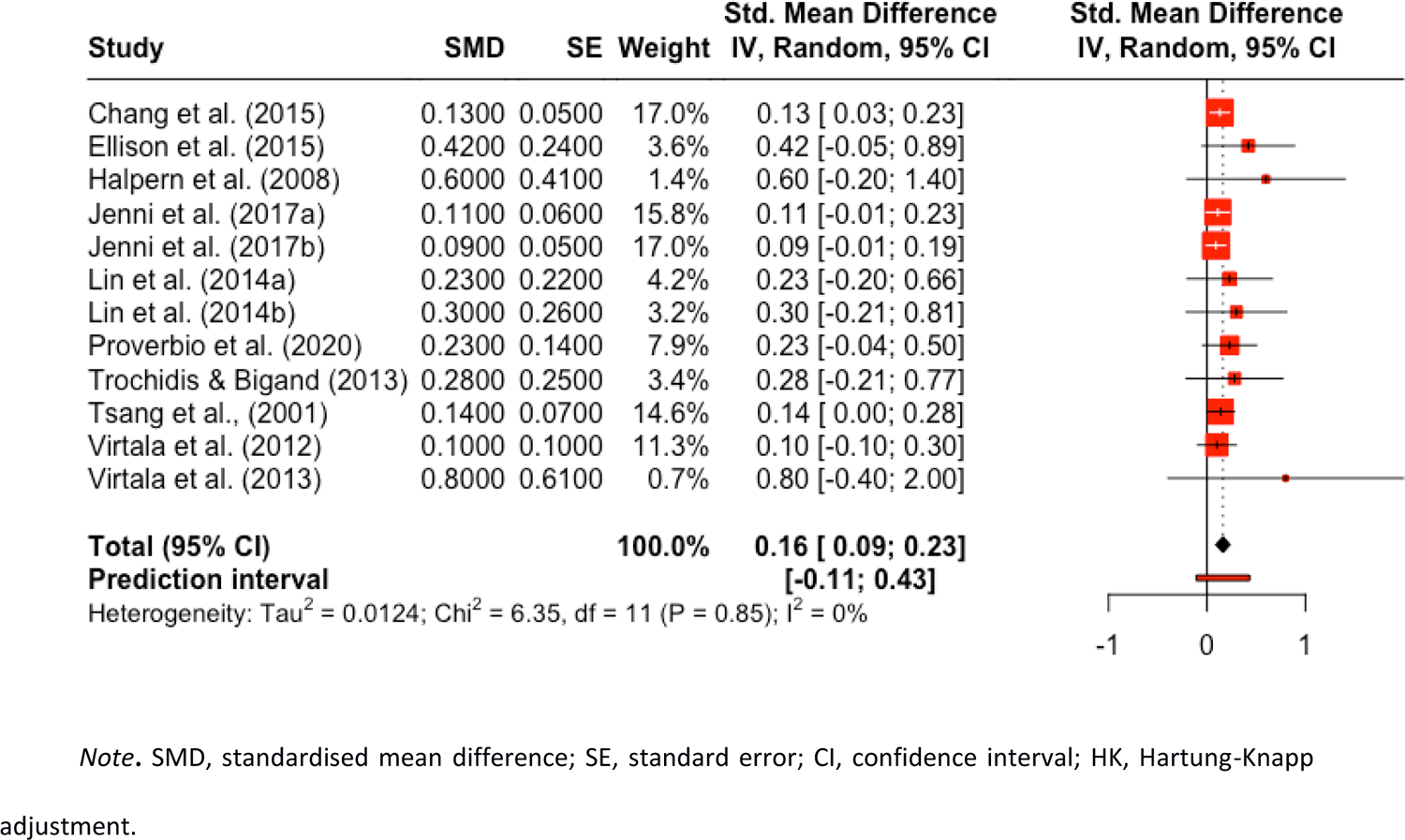
EEG Meta-analysis forest plot

### Meta-analysis of Neuroimaging Studies

#### ALE Meta-analyses

The ALE meta-analysis of major-minor (**Table 3** & **Figure 4**) revealed significant convergence in (1) left superior temporal gyrus Brodmann area 22 (STG-L BA22a), (2) right medial frontal gyrus (MedFG-R BA6), (3) right transverse temporal gyrus (TTG-R BA41), (4) right cingulate gyrus (CG-R BA31), and (5) right caudate (CAU-R). The ALE meta-analysis of minor-major revealed significant convergence in the (1) left superior temporal gyrus (STG-L BA22b) and (2) right superior temporal gyrus (STG-R BA22).

**Figure 4.**
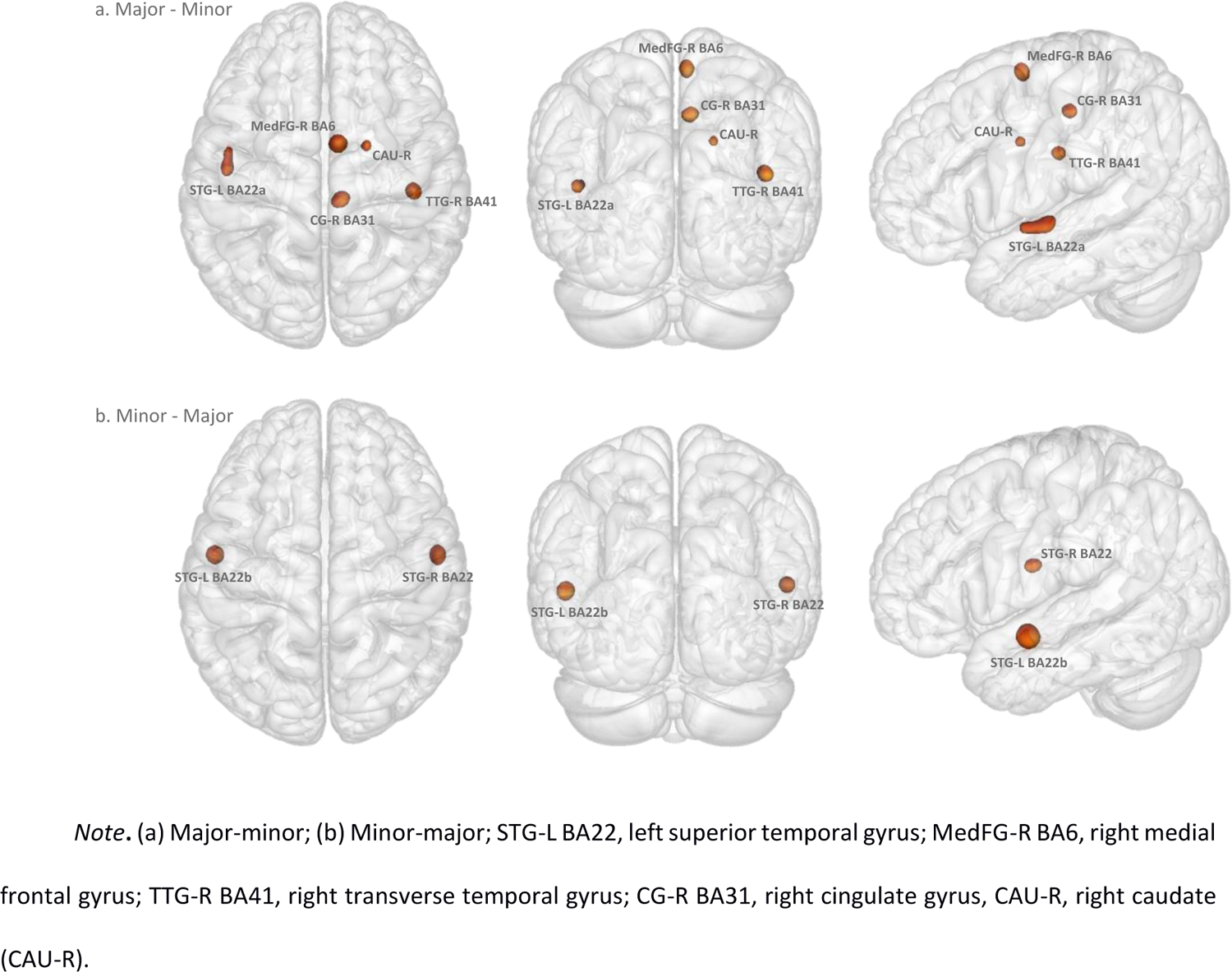
ALE Meta-analysis

**Table 3.** ALE Meta-analytic Results of fMRI Studies Comparing Major vs Minor Mode at Cluster Level Inference p < .005 (FWE)

| Cluster number | Volume (mm<sup>3</sup>) | MNI coordinates |  |  | ALE | P | Z | Label (Side region BA) |
| --- | --- | --- | --- | --- | --- | --- | --- | --- |
|  |  | x | y | z |  |  |  |  |
| 1. Major – Minor: 65 foci, 7 experiments, 119 subjects |  |  |  |  |  |  |  |  |
| 1 | 1272 | -52 | -14 | 0 | 2E-02 | 2E-07 | 5.1 | L Superior Temporal Gyrus BA22 |
| 2 | 824 | 6 | -4 | 60 | 2E-02 | 2E-08 | 5.5 | R Medial Frontal Gyrus BA6 |
| 3 | 808 | 46 | -24 | 10 | 2E-02 | 8E-09 | 5.7 | R Transverse Temporal Gyrus BA41 |
| 4 | 768 | 8 | -28 | 40 | 2E-02 | 3E-08 | 5.4 | R Cingulate Gyrus BA31 |
| 5 | 488 | 20 | -2 | 22 | 2E-02 | 7E-07 | 4.8 | R Caudate |
| 2. Minor – Mayor: 33 foci, 8 experiments, 138 subjects |  |  |  |  |  |  |  |  |
| 1 | 1096 | -60 | -8 | -6 | 2E-02 | 1E-10 | 6.4 | L Superior Temporal Gyrus BA22 |
| 2 | 1040 | 60 | -8 | -2 | 2E-02 | 5E-09 | 5.7 | R Superior Temporal Gyrus BA22 |

ALE, anatomic likelihood estimation; BA, Brodmann area; FWE, family-wise error correction; P, p-value; Z, peak z-value; R, right; L, left.

The functional characterization of each ROI focused on behavioural domains (e.g., action, perception, emotion, cognition and interoception) and paradigm classes (e.g., pitch discrimination, finger tapping, music comprehension, and music production) included in the BrainMap platform (**Table S5**). MACM was performed to functionally segregate the behavioural contribution and the patterns of co-activation (or FC) of music-mode-related region-of-interest (ROI) resulting from the major-minor ALE (n=5) and minor-major ALE (n=2) meta-analyses (**Table S6**).

### Major – Minor MACM

STG-L BA22a showed significant FC with right superior temporal gyrus BA22, left thalamus, and right cingulate gyrus BA32. Relevant behavioural domains within its boundaries include execution, speech, language memory, music, emotion, and auditory processing; and experimental paradigms including finger tapping, music comprehension and production, and oddball, phonological, pitch, semantic, and tone discrimination.

MedFG-R BA6 showed significant FC with right thalamus, left and right cerebellum, and left insula BA13. Relevant behavioural domains include execution, speech, memory, music, emotion, reward, and auditory processing; and experimental paradigms including emotion induction, finger tapping, music comprehension and production, and oddball, phonological, pitch, semantic, and tone discrimination.

TTG-R BA41 showed significant FC with left superior temporal gyrus BA41, right precentral gyrus BA6, and left and right thalamus. Relevant behavioural domains include execution, speech, memory, music, emotion, reward, and auditory processing; and experimental paradigms including finger tapping, music comprehension and production, and phonological, pitch, semantic, syntactic, and tone discrimination.

CG-R BA31 showed significant FC with surrounding areas within the cingulate gyrus. Relevant behavioural domains include execution, speech, memory, music, emotion, reward, and auditory processing; and experimental paradigms including finger tapping, music comprehension, and music production.

CAU-R showed significant FC with left superior frontal gyrus BA6 and left claustrum. Relevant behavioural domains include execution, memory music, emotion, reward, and auditory processing; and experimental paradigms including finger tapping, music comprehension, and oddball and tone discrimination.

### Minor – Major MACM

STG-L BA22b showed significant FC with right superior temporal gyrus and left precentral gyrus. Relevant behavioural domains include execution, speech, memory, music, emotion, reward, valence, and auditory processing; and experimental paradigms including emotion induction, finger tapping, music comprehension, and phonological, pitch, semantic, syntactic, and tone discrimination.

STG-R BA22 showed significant FC with left superior temporal gyrus, medial frontal gyrus BA6, left middle frontal gyrus BA9, and left insula BA13. Relevant behavioural domains include execution, speech, memory music, emotion, reward, valence, and auditory processing; and experimental paradigms including emotion induction, finger tapping, music comprehension, and phonological, pitch, semantic, syntactic, and tone discrimination.

## Discussion

The major-minor dichotomy is a fundamental aspect of Western tonal music, and it refers to the two most commonly used modes, or scales. The major mode is characterised by a bright, happy, and uplifting sound, while the minor mode is characterised by a darker, sadder, melancholier, and introspective sound. For this comprehensive review and meta-analysis, we conducted a thorough literature search and selection process that identified 69 relevant studies on the perception of musical mode (54% mode-centred) that included 4,449 participants. While most studies focused on major-minor mode, some studies investigated emotional responses to music, with musical mode being the independent variable, and the emotional response being the dependent variable. The geographical distribution of the studies was mainly in the Western regions; however, the USA, Canada, Finland, and Japan had the most significant number of studies. Also, most of the research was published after 2010. Furthermore, our review shows that behavioural studies represent most articles, with a few applying exclusively physiological measures and others using both. It is also worth noting that the type of musical stimulation used in the studies varied, with some using short fragments of music while others used longer pieces. In terms of the quality assessment, 90% of the studies were scored as high-quality publications, while only one study was scored as low quality.

Overall, our review suggests that the major-minor dichotomy has been a significant focus of research in music perception, although it is often investigated alongside other features of music. It is important to consider the role of musical mode in investigations of emotional responses to music, as this dichotomy can be a powerful inducer of happy/sad emotions, and thus, has potential clinical applications for mood disorders, for example. Here, we propose a framework to describe a *Major-Minor Mode(l)* in music that can be used to provide insights into the ways in which music affects our emotions, cognition, and physiology.

### Affective Evaluation of the Major-Minor Dichotomy

The affective evaluation of major and minor modes can be seen in various musical compositions. For example, many joyful and celebratory songs are written in major keys, such as “Happy Birthday” or “The Star-Spangled Banner.” These songs use major chords and melodies that are uplifting and energetic, which evoke feelings of joy, excitement, optimism, and celebration (Pilcher et al., 2014). On the other hand, minor keys are often used in music to express sadness, grief, or melancholy. For instance, many funeral dirges or requiems are composed in minor keys, such as the famous *Funeral March* by F. Chopin or W.A. Mozart’s *Requiem in D Minor*. These pieces use minor chords and melodies that are haunting, sorrowful, and poignant, which evoke feelings of sadness, introspection, and longing.

However, the affective evaluation of major and minor modes is not always straightforward. While major keys are typically associated with happiness, they can also be used to express other emotions, such as anger or sarcasm. For example, *The Entertainer* by S. Joplin is a lively ragtime piece written in a major key that sounds cheerful and upbeat, but the composer intended it to be ironic and satirical. Similarly, while minor keys are often associated with sadness, they can also convey a sense of mystery, suspense, or even romance. For example, the *Moonlight Sonata* by L. van Beethoven is a well-known composition written in a minor key that expresses a sense of introspection and longing, but also contains moments of beauty and tranquillity.

### The Valence-Arousal Model

Music is a complex and powerful medium that can evoke a wide range of emotions and experiences in listeners. However, the link between music and emotions can be intricate and not always apparent. The same piece of music can elicit different emotional responses in different people, and individual differences such as personality, musical preference, and cultural background can all play a role in shaping listeners’ emotional experiences, as it will be discussed later. One way that researchers have tried to understand the emotional impact of music is through the so-called valence-arousal model.

Whilst the association between musical modes and distinct emotional responses dates back to the Greek theorists and has been debated along the whole history of music (Wiering, 1998), the first scientific report that highlighted the affective connotation of musical modes was by Hevner (1937), who asked listeners to classify musical excerpts with “sad” or “happy” adjectives. The results indicated that major mode was associated with “happy” adjectives whilst the participants selected more “sad” adjectives when the music was played in minor mode. The affective dichotomy major/happy minor/sad has been confirmed over the following decades, and other specific emotions have been studied related to this dichotomy. For example, the major mode has been associated with an experience of joy, serenity and vitality, whereas minor mode tends to be perceived as nostalgic, subdued, dark and wistful (Bonetti & Costa, 2019; Brattico et al., 2011; Crowder, 1985; Peretz et al., 1998).

At its core, the valence-arousal model suggests that emotions can be described using two primary dimensions: valence and arousal. Valence refers to whether an emotion is positive (e.g., happy, content) or negative (e.g., sad, angry) (Koelsch et al., 2006). Arousal, on the other hand, refers to the intensity or activation level of an emotion, ranging from low arousal (e.g., calm, relaxed) to high arousal (e.g., excited, energised) (Reisenzein, 1994). The valence-arousal model has been widely applied in music research, with studies using it to explore how different types of music affect listeners’ emotional experiences (Chang et al., 2015; McConnell & Shore, 2011; van der Zwaag et al., 2011).

### Pleasure and Reward

The major-minor dichotomy has been explored in various studies to understand the relationship between music, pleasure, and reward. In terms of musical pleasure, research has shown that people tend to prefer music in major mode, given the positive psychological character generally associated with it (Blood & Zatorre, 2001; Dobrota & Ervegovac 2015; McConnell & Shore, 2011). One influential theory is that listeners experience a sense of reward or pleasure when they are able to anticipate and predict the harmonic structure of a musical piece (Huron, 2006; Koelsch, 2014; Pando-Naude et al., 2021; Pearce, 2005). According to this theory, the major-minor dichotomy is particularly important because it creates expectations about how a piece of music will unfold. When listeners can accurately predict the next harmonic change, they experience a sense of satisfaction and pleasure. In Hunter et al. (2008), for instance, participants were asked to rate musical excerpts using scales for happiness/sadness and pleasantness/unpleasantness, with the results showing that listeners found music they rated as happy also pleasant and, vice versa, sad music tended to be considered unpleasant. However, minor mode is not exclusively related to unpleasant judgments. Indeed, in Green et al. (2008), participants did not provide higher ratings of liking for major over minor melodies, although they perceived major music happier than minor. According to Vuoskoski & Eerola (2012), two personality traits, openness to experience and empathy, are associated with liking sad music. Similarly, Bonetti & Costa (2016) found that the preference for short melodies in minor mode is positively related to the level of fluid intelligence and openness to experience. Importantly, the use and preference for major and minor modes can be influenced by a wide range of variables, including individual strategies of emotional regulation through music, as well as potential clinical conditions that may affect its perception, as described in the next paragraphs.

### On the Origins and Universality of Major-Minor Affective Connotation

One of the most intriguing questions related to the study of major and minor modes concerns the potential generalizability of the typical happy-sad emotional response beyond the Western tonal context. So far, all the mentioned studies investigated Western music and Western listeners, nevertheless, other studies found some commonalities in music-induced emotions also among people of other cultures. Specifically, both Hoshino (1996) and Fang et al. (2017) found that excerpts in major mode induced pleasure and happiness, whereas the ones in minor mode were associated with greater tension among Japanese and Chinese people, respectively, consistently with studies on Western listeners. This evidence seems to suggest that the affective connotation of musical mode might rely on a universal dimension. However, a recent cross-cultural study conducted between Australia and Papua New Guinea, reported that major mode does not induce greater happiness than minor in a group of people with minimal exposure to Western music (Smit et al., 2022). This remarks that the typical major/happy minor/sad association might partly be a culturally learned association, although the debate is still open, and more research is needed.

To understand the influence of cultural factors in the development of the major-happy and minor-sad association, researchers have explored the psychoacoustic characteristics that might underlie the development of this typical dichotomous connotation. For instance, Bowling et al. (2010) proposed that the emotional connotation evoked by different tonal relationships in music (such as major and minor mode), may be related to the emotional information conveyed by the quality of human speech. Specifically, the spectral characteristics of excited speech tend to align more closely with the spectral characteristics of intervals found in major music. Conversely, the spectral characteristics of subdued speech tend to align more closely with the spectral characteristics of intervals that are distinctive to minor music. These results are in line with Cook et al. (2006) who suggested that the harmonic relationships of minor mode music are similar to sad speech prosody. Also, processing of emotional vocalizations and music seems to involve common neural mechanisms (Proverbio et al., 2020). This line of research emphasizes thus the similar acoustic cues present in music and vocalizations to express emotions, determining music’s ability to convey emotions in a universal manner. However, several cross-cultural studies indicate that cultural exposure and socialization play a role in shaping the perception of emotional vocalizations (for a meta-analysis see Laukka & Elfenbein, 2021). Sauter et al. (2010) found that while certain primarily negative emotions have vocalizations that can be recognized across cultures, most positive emotions are communicated with culture-specific signals. Similarly, Gendron and colleagues (2014) proposed that the associations between particular vocalizations and specific perceived mental states might not universally exist across different cultures. These results do not rule out the possibility that the affective connotation of major and minor modes may stem from similar patterns of emotional vocalization. However, universality remains a hypothesis not entirely supported.

### Beyond Happiness and Sadness: the Complexity of the Affective Experience of Music

Recognizing the importance of complexity in music is fundamental to our appreciation of this extraordinary art form. Music has indeed the unique ability to elicit in the listener a spectrum of emotions, ranging from the simplest to the most complex and profound, such as aesthetic emotions (Brattico & Pearce, 2013). An aesthetic experience is defined as a special state of mind, in which focused attention plays a pivotal role, and which is responsive not to bodily needs (such as appetitive and mating functions) but rather offers pleasures for the mind (Marković, 2012). Aesthetic emotions are not simply a subset of everyday emotions, but rather a distinct category of emotions that are elicited by the unique features of art, such as form, color, and composition (Pelowski et al., 2017). The affective connotation of major and minor modes, in this regard, plays a fundamental role. In fact, the major-minor dichotomy is generally utilized in music to evoke discrete emotions of happiness and sadness, respectively. Nevertheless, combinations of music in major and minor mode, along with the context and other listener-related elements, such as their attitude or mood, can engender more nuanced aesthetic responses. For instance, skilful exploitations of major and minor mode in music, such as transitioning from a minor mode to a major mode can create a powerful emotional impact. An example is Beethoven’s *Symphony No. 5*, where the symphony begins in C minor but in the final movement it resolves triumphantly into C major, conveying a sense of victory and triumph. Similarly, Tchaikovsky’s *Symphony No. 5* starts in B minor, evoking a sense of sadness and tension, but in the last movement, it shifts to B major, creating a moment of resolution and emotional release. Accordingly, studies employing naturalistic paradigms seem more appropriate in capturing the complex “aesthetic” and mixed affective responses induced by music, reaching far beyond the basic categories (e.g., happiness vs. sadness) (Brattico & Alluri, 2022; Trost et al., 2012).

The temporal evolution of aesthetic emotions in music has been well-addressed in a study by Brattico et al. (2013), where the authors suggested that the aesthetic experience of music occurs on distinct temporal and spatial levels, highlighting a difference between aesthetic judgment and conscious liking. The authors propounded a dynamical description of the aesthetic experience of music suggesting that the complete actualization of a musical aesthetic experience requires a particular (aesthetic) attitude, intentionality, attention, and the appropriate context. Specifically, music can express and induce basic and discrete emotions, such as happiness and sadness, conveyed for example by major-minor dichotomy, a sort of early emotional reaction. Eventually, the combination of cognitive processes, attentional mechanisms, and contextual factors is what leads to outcomes of aesthetic emotions, aesthetic judgments, and attitudes.

Furthermore, the familiarity with music plays a significant role in shaping emotional responses. Familiarity can influence listeners’ perceptions of what constitutes an optimal level of complexity and the resulting aesthetic experience (Mechner, 2018; Schellenberg et al., 2008a; Verhaeghen, 2018). In the perspective of understanding the optimal complexity in music, researchers often refer to the Goldilocks Effect, namely the phenomenon where music that is neither too predictable nor too unpredictable is preferred, even in infancy (Kidd et al., 2012). Accordingly, familiarity with certain musical structures and patterns can enhance listener’s emotional engagement and enjoyment, by influencing listeners’ perceptions of what constitutes an ideal musical experience.

The emphasis on listener’s expectations and attitude is also in line with the concept of affective forecasting, namely a psychological phenomenon that involves predicting and anticipating one’s emotional reactions to future events or experiences (Wilson & Gilbert, 2003). Musical enjoyment and preference may be thus closely related to the contingency, the context, and the individual. Indeed, according to the so-called arousal-based goals (North & Hargreaves, 2000), people tend to prefer musical pieces deemed appropriate to the context. For this reason, musical pieces with different arousal and valence are usually selected for relaxing rather than for a sport activity. This also contributes to explain the paradoxical preference for negative music (i.e., minor-mode) in certain contexts, along with the effect of individual differences (e.g., Bonetti & Costa, 2016; Vuoskoski & Eerola, 2012).

In sum, comprehending the emotional impact of the major-minor dichotomy in music is fundamental, considering its centrality, especially in Western music. Simultaneously, embracing the complexity in music necessitates acknowledging the multifaceted nature of emotional and aesthetic elements. In this regard, the affective experience associated with music appears nuanced, intricate, and, as a result, cannot be dissociated from the consideration of factors linked to the individual and the context. Moreover, the familiarity with music and the Goldilocks Effect contribute to shaping emotional responses, enhancing the listener’s engagement and enjoyment of the musical experience.

### Association between Type of Stimulation and Elicited Behavioural and Physiological Responses

The reviewed studies aimed to investigate the perception of musical mode using a variety of stimuli such as chords, excerpts, musical pieces, and tone scrambles (**Table 1**). The major mode was perceived as relatively happy and bright, whereas music in minor mode tends to be perceived as dark and sad (Hevner, 1937; Kastner & Crowder, 1990). Isolated chords and very short melodies prove to represent sufficient stimulation to elicit these distinct affective responses (Bonetti & Costa, 2019; Ellison et al., 2015; Pallesen et al., 2003). Consistently, major and minor chords evoke differential responses both on a physiological and neural level regardless of whether they are presented in isolation(Pallesen et al., 2005), inserted in an oddball paradigm (Putkinen et al., 2014; Virtala et al., 2011, 2013), or in a sequence (Suzuki et al., 2008). Moreover, musical expertise is partly reflected in terms of perceptual differences of major-minor chords and its implications will be further discussed (Pallesen et al., 2015; Putkinen et al., 2014; Virtala et al., 2012).

### Behavioural

Regardless of stimulus duration, in studies employing pure tones, listener’s ability to discriminate between major and minor modes appears lower, resulting in a reduced perception of the characteristic major/happy and minor/sad association (Chubb et al., 2013; Dean & Chubb, 2017; Mednicoff et al., 2018). Most of the reviewed studies (70%) used longer-lasting stimuli, and in the bulk of them, musical excerpts and melodies were selected based on previous music research. In a few studies, the excerpts/melodies were composed for the purpose (Bonetti & Costa, 2019; Franco et al., 2017; Gosselin, 2005; Halpern, 1984; Trochidis & Bigand, 2013), a customised selection of real songs/excerpts (Al’tman et al., 2000; Gregory et al., 1996; Hunter et al., 2008; Schellenberg et al., 2008; Tai & Kuo, 2019), and partly even chosen by the participants (Brattico et al., 2016). Presumably, a longer duration increases the possible effects arising from other musical features besides mode, such as tempo, rhythm, or timbre, resulting in independent variables difficult to control for. Indeed, different durations of real-life musical stimuli may also be associated with different levels of arousal elicited by major-minor modes (Fang et al., 2017; Kastner & Crowder, 1990; van der Zwaag et al., 2011). However, similar stimuli do not necessarily entail similar results, mainly due to the different methodological tools used to assess the elicited responses (Khalfa et al., 2005; Lin et al., 2014; Trochidis & Bigand, 2013).

Remarkably, the sensory-perceptual discrimination between major and minor chords, as measured by the MMN paradigm, develops very early in infants (Virtala et al., 2013), whereas the ability to explicitly associate emotions to the two modes emerges later, starting from 3-4 years (Kastner & Crowder, 1990), and is fully developed at the age of 8-9 years (Nieminen et al., 2012). The development of this ability appears to consistently correlate with the development of fluid intelligence and cognitive maturation (Bonetti & Costa, 2016).

The behavioural meta-analysis conducted in this study sought to investigate the impact of major and minor music modes on emotional connotations in a behavioural context. By synthesizing data from 12 experiments, we aimed to provide a comprehensive understanding on how musical mode affects emotional responses. Our results suggest that musical mode significantly impact emotional responses, as evidence by the positive SMD estimate. However, the substantial heterogeneity among the included studies underscores the need for further exploration of moderators and potential sources of variation in the relationship between musical mode and emotion.

### Autonomic

Several studies have investigated the effect of musical mode on emotional autonomic responses using physiological measures such as skin conductance level (SCL), heart rate variability (HRV), and salivary cortisol levels. Van der Zwaag et al. (2011) found that musical pieces in minor mode elicited higher arousal compared to the major mode, suggesting that it is due to the higher degree of uncertainty and surprise arising from minor mode. However, the results of Kastner & Crowder (1990) contradict this finding, with the major mode eliciting greater arousal. The discrepancy between these studies may be attributed to different reasons, for example, the stimuli duration and/or the age of the participants. Suda et al. (2008) found that major mode music had a greater effect on reducing stress, as evidenced by a significant decrease in salivary cortisol levels compared to minor mode music and a control group without auditory stimulation. This result was supported by the activation of the dorsolateral prefrontal cortex, strongly associated with positive mood, in response to major mode music using optical topography (OT). This evidence is consistent with previous studies which indicated that the activation of the dorsolateral prefrontal cortex is strongly associated with a positive mood (Habel et al., 2005; Markowitsch et al., 2003).

### Neural Correlates

The neural correlates of musical mode have been investigated through various neuroimaging techniques, but the results have not always been consistent. For example, EEG recordings by Davidson (1988) suggested that greater activity in the left frontal lobe is associated with positive emotions like happiness, while the right frontal area is related to negative emotions like sadness and fear. However, an fMRI study by Khalfa et al., (2005) did not confirm this lateralization, as participants displayed stronger left activation when exposed to sad musical excerpts in minor mode and slow tempo. Specifically, the minor versus major mode contrast showed greater activity in the posterior cingulate cortex, left orbital and mid-dorsolateral prefrontal cortex, and limbic structures. These results were supported by other fMRI studies (Green et al., 2008; Pouladi et al., 2011). EEG studies have also confirmed the involvement of the medial frontal and right sensorimotor activations in musical mode processing (Lin et al., 2014).

Trochidis & Bigand (2013) conducted an EEG experiment to investigate emotional responses to different musical pieces played in major, minor, and Locrian mode. Participants rated major mode pieces as happier and more serene compared to minor and Locrian modes. In EEG frontal activity, major mode was associated with increased alpha activation in the left hemisphere compared to minor and Locrian modes, which increased activation in the right hemisphere, supporting the valence lateralization model suggested by Davidson (1988). It should be noted, however, that in these studies, stimuli presented to the subjects were not specifically composed with well controlled mode for interpreting the brain responses. Thus, the presence of variable musical parameters may have caused confounding effects in the neuroimaging data, partly explaining the discordant results.

In contrast, Suzuki et al. (2008) employed only major-minor chords in a PET study to investigate the neural correlates of major-minor modes when both are perceived as pleasurable. Results indicated that minor chords perceived as beautiful strongly activated the right striatum, associated with reward and emotion processing, while “beautiful” major chords elicited a higher activity in the left middle temporal gyrus, an area involved in cognitive processes such as orderly information and semantic memory processing (Gernsbacher & Kaschak, 2003; Onitsuka et al., 2004).

The involvement of the motivational-reward circuit in music-induced pleasure is consistent with previous findings (Blood & Zatorre, 2001) and mainly concerns minor mode music, as reported by Green et al. (2008), Pallesen et al. (2005), and Pouladi et al. (2011). However, the investigation of brain responses to major and minor music yields very discordant evidence. Music in minor mode has been associated with greater brain activation than major music in both fMRI (Green et al., 2008; Nemoto et al., 2010), EEG (Proverbio et al., 2020), and MEG-based studies (Jomori et al., 2013). On the other hand, several studies reported increased activation elicited by major music compared with equivalent minor (Chang et al., 2015; Jeong et al., 2011; Tsang et al., 2001). Interestingly, two studies using event-related fMRI, failed to find brain regions differentially activated by listening to music in major vs. minor mode, when presented as triad sequences (Mizuno & Sugishita, 2007), or melodic sequences (Lee et al., 2011). Similar results were reported by Maslennikova et al. (2013), showing differences in evoked EEG responses substantially based on perception of consonant and dissonant chords rather than musical mode.

Considering these divergent findings, it might be conjectured that the neural correlates of major-minor perception are a consequence of the subjective perceived emotional connotation and pleasantness rather than major and minor mode itself. This is in line with Tabei (2015) who highlighted that the emotional expression of music and emotional responses evoked in the listener are distinctly processed in the brain.

In addition to the fundamental differences between the neuroimaging methods described, the discrepancies in literature might be ascribed to the overlapping of some of the neural correlates related to happiness and sadness perceived in music (Brattico et al., 2011; Khalfa et al., 2005; Mitterschiffthaler et al., 2007). Brain structures such as the amygdala, hippocampus, striatum, and regions of the reward circuit are involved in both pleasantness and unpleasantness perception in music (Blood et al., 1999; Blood & Zatorre, 2001; Suzuki et al., 2008). Thus, the overlap in neural correlates between happiness and sadness in music perception can make it challenging to isolate the specific neural correlates associated with the major-minor mode dichotomy. This is because brain structures such as the amygdala, hippocampus, striatum, and regions of the reward circuit are involved in both pleasantness and unpleasantness perception in music.

For example, a study by Blood & Zatorre (2001) found that listening to pleasant music activates the striatum, a brain region associated with reward and motivation, whereas listening to unpleasant music activates the amygdala, a brain region associated with negative emotions such as fear and sadness. However, these regions are also activated in response to other stimuli (Kim & Hamann, 2007), making it difficult to determine the specific contribution of mode to these neural responses.

Therefore, while there is evidence to suggest that major and minor modes elicit different emotional responses in listeners, the specific neural correlates associated with these responses are still not fully understood and are likely to be influenced by a range of subjective and individual factors (Eerola & Vuoskoski, 2013).

The EEG meta-analysis conducted in this study synthesized findings from twelve independent experiments to investigate the impact of music mode (major vs minor) on EEG activity as measured by the SMD. This effect size, which falls within the moderate range according to conventional guidelines, suggest that the presence of music mode is associated with a meaningful change in EEG, presumably driven by emotional connotations. Notably, the low heterogeneity observed in the results suggests that the effect of music mode on EEG activity was relatively consistent across the different studies.

The fMRI ALE meta-analysis revealed significant convergence of areas that are more likely to be active by major in comparison with minor mode including auditory, premotor, limbic, and basal ganglia. STG-L BA22a is associated with a wide range of functions including language processing, memory, music perception, and emotion. Its functional connectivity with the right superior temporal gyrus suggests an interhemispheric relationship in processing major-minor distinctions in music. MedFG-R BA6 is linked to executive functions, speech, memory, and emotion. The connectivity with the thalamus and cerebellum points to its role in motor coordination and emotional processing during musical activities. TTG-R BA41 is involved in auditory processing and speech perception. Its connectivity with the left superior temporal gyrus (BA41), right precentral gyrus (BA6), and thalamus suggests its role in the discrimination of pitch and phonological aspects of music. CG-R BA31 is functionally connected with surrounding areas within the cingulate gyrus and is linked to various cognitive and emotional processes. It likely contributes to emotional and cognitive aspects of music. CAU-R is an area of the basal ganglia associated with reward processing and motor functions. Its connectivity with the superior frontal gyrus (BA6) and claustrum suggests involvement in motor coordination, memory, and emotion processing during musical activities.

In contrast, areas more likely to be active by minor mode include areas within the auditory cortex. STG-L BA22b demonstrated significant FC with the right superior temporal gyrus and the left precentral gyrus, suggesting an intricate interplay between these regions in processing musical stimuli, especially those related to the discrimination of minor and major modes. Importantly, the behavioural domains encompass a wide range of cognitive functions, including execution, speech, memory, music perception, emotion processing, reward mechanisms, valence appraisal, and auditory processing. This multifaceted engagement implies that STG-L BA22b plays a versatile role in the perceptual and emotional aspects of music, contributing to both the analysis and emotional interpretation of musical content. Additionally, STG-R BA22 exhibited functional connectivity with several brain regions, including the left superior temporal gyrus, medial frontal gyrus BA6, left middle frontal gyrus BA9, and left insula BA13. These connections implicate a broad array of cognitive and emotional functions, mirroring those associated with STG-L BA22b, including execution, speech, memory, music perception, emotion processing, reward processing, valence assessment, and auditory processing. This extensive involvement suggests that STG-R BA22 is engaged in the comprehensive processing of musical content, from the analysis of pitch and syntax to the emotional response evoked by music. Furthermore, the presence of STG-R BA22 in experimental paradigms related to emotion induction, finger tapping, and various discrimination tasks reinforces its role in the intricate perception and interpretation of musical elements.

Together, these meta-analytic results revealed significant alterations in EEG activity and distinct neural networks associated with major and minor modes, shedding light on the consistent and multifaceted impact of music on the human brain.

### Individual Factors in the Perception of the Major-Minor Dichotomy

#### Individual Differences in Major-Minor Discrimination

In some studies, researchers have found that there are differences in how individuals discriminate between major and minor musical stimuli. However, the effect sizes are relatively modest, meaning the differences are not very significant across individuals. Recently, Chubb et al. (2013) and Dean & Chubb (2017) reported that most listeners (70%) performed near chance when asked to classify rapid tone-scrambles composed of multiple copies of notes in G-major vs G-minor triads as major or minor, whereas the remaining listeners performed nearly perfectly. Interestingly, these results were replicated when the stimuli are slowed down (Mednicoff et al., 2018).

Moreover, Adler et al. (2020) demonstrated that, by employing the same stimuli, a similar bimodal distribution in 6-month-old infants. However, these latter studies have been carried out by using tone-scrambles, which are random sequences of pure tones, instead of more natural harmonic tones, typically employed in the investigation of major-minor perception. Indeed, in contrast, several studies suggest that nearly all listeners are able to sense some sort of mode-related differences when exposed to more prototypical and realistic sounds (Bonetti & Costa, 2019; Pallesen et al., 2003; Temperley & Tan, 2013; Virtala et al., 2011), including non-Western listeners (Fang et al., 2017; Hoshino, 1996), and newborn infants (Virtala et al., 2013).

#### Effect of Musical Expertise on Major-Minor Perception

As reported in **Table 2**, nearly half of the studies reviewed (n=30) investigated the effect of musical expertise on music mode perception by comparing musicians and non-musicians. Halpern et al. (2008), for instance, created pairs of identical melodies except for the mode, and both musicians and non-musicians were asked to judge the melodies as major vs. minor, or happy vs. sad. The main aim of this experiment was to analyse the processing of music at the point in time of the first note that differentiated major or minor mode (critical note). The ERP showed a large N1 and P2 component in both musicians and non-musicians to the onset of the critical note, but, interestingly, the musicians showed a late P3 (LPC) to the critical note only for minor melodies in both tasks. Non-musicians, in contrast, showed a dissociation between their behavioural and physiological responses, namely, they could easily classify the melodies as happy or sad, but they did not show any LPC response. The late P3 recorded in musicians only for minor mode stimuli is consistent with the fMRI study by Khalfa et al. (2005), indicating a specific neural activity only in response to minor tunes. This tendency might be explained by the fact that minor-mode music is less common than major-mode music. Indeed, Huron (2006) found that when musicians were asked to think about a random chord, 94% of them imagined a major mode chord. Hence, the major mode appears to be the most “expected” mode, and the minor mode could elicit a higher brain response as an infrequent, deviant stimulus (Jenni et al., 2017; Trochidis & Bigand, 2013). Similarly, major music is deemed more “correct” than minor music (Ellison et al., 2015). Nonetheless, the modulation of musical mode perception by expertise was not systematically observed in all the studies. For instance, Pallesen et al. (2005) did not find any consistent difference between musicians and non-musicians when they listened to major, minor, and dissonant isolated chords.

To clarify the underlying mechanisms of musical mode and their development based on musical expertise, previous studies have investigated the distinct frequency-bands connecting neuronal networks, which are involved in music processing. Lower frequency bands, such as theta and alpha, are considered to be processing “cognitive” information, whereas higher frequency bands (beta and gamma) are involved in processing more basic features (von Stein et al., 2000). Pallesen et al. (2015) reported a stronger gamma-band activity in response to non-prototypical chords (i.e., dissonant and mistuned) in musicians, while beta-band activity was stronger in non-musicians listening to minor chords. Additionally, a general reduction of gamma-band activity in response to major chords was observed in both groups. This effect may reflect ‘easy’ processing of this mode due to its frequent occurrence in Western music. Hence, high-frequency EEG bands appear to be crucial in processing more complex musical features, and the activity may be modulated by musical expertise. This view is consistent with the findings of a recent study by Jenni et al. (2017) who reported that minor and major compositions distinctively modulated synchronisation of neuronal activities in high-frequency ranges (beta and gamma) in frontal regions, with increased activity in response to minor compositions especially in musicians and experts. This suggests that high-frequency EEG activities also carry musical mode information in addition to highlighting the role of musical expertise in enhancing the efficient processing and sensitivity to mode. Also, the involvement of frontal regions and limbic structures in music mode processing is consistent with previous studies (Green et al., 2008; Khalfa et al., 2005; Pallesen et al., 2005).

In terms of early musical training, previous studies suggest that minor mode chords can elicit an automatic error response in the brain (MMR) even in infants (Virtala et al., 2013), whereas similar stimuli do not evoke any significant error response (MMN) in older children without musical training (Virtala et al., 2012). This inconsistency might indicate a sort of predisposition towards the major-minor neural discrimination that tends to decline during the development when musical training is absent. However, the utilisation of more complex paradigms may help to level these differences, since Tervaniemi et al. (2014) found prominent MMN responses to sound changes in both musicians and non-musicians by using the melodic multi-feature paradigm. This paradigm is faster and has a higher ecological validity than previous MMN paradigms.

Finally, musical expertise does not determine only differences in terms of neural response to mode, but it also affects individual preference. In fact, well-trained musicians consider musical excerpts in minor mode as more beautiful and pleasant than non-musicians (Brattico et al., 2016).

#### Effect of Age on Major-Minor Perception

When investigating the behavioural and neural correlates of musical mode, a fascinating question is whether musical mode perception changes throughout life. Dalla Bella et al. (2001) reported that 5-year-old children could discriminate between happy and sad musical excerpts, but would only describe the excerpts in terms of tempo, whereas from the ages of 6-8 they utilised both tempo and mode, like adults. Similar results were obtained when children aged 3-4 and 7-8 were asked to associate a happy or sad face to music in major and minor mode. Moreover, 7-8-year-old children responded significantly differently to popular tunes in major and minor mode, but 3-to-4-year-old children showed almost no difference (Gregory et al., 1996). Both studies suggest that the emotional perception related to musical mode develops between the ages of 4 and 7.

However, the beginning of the development of the major/happy and minor/sad association around the age of 4 is not supported by all the studies reviewed here. For example, Kastner & Crowder (1990) found that 3-year-old children show the ability to match schematic faces (such as sad or happy) to musical excerpts based on the musical mode. More recently, an experiment by Franco et al. (2017), in which 3–to-6-year-old children were asked to match facial expressions to the music tracks heard, showed that even the youngest children were able to correctly identify the intended emotion in music, whereby the recognition of happy music appeared facilitated in the instrumental condition. To sum up, most studies reviewed suggest that the affective connotation of major and minor mode is individually developed around the age of 4 and it is entirely acquired around the age of 7-to-8.

Recently, in a cross-modal association study, Bonetti & Costa (2019) found that the association between major mode and happiness, and minor mode and sadness when infants and adults were exposed to musical chords increased from 58% at the age of 4 to 61% at the age of 5, 72% at the age of 6, reaching 92% in adults. Similarly, another study found that 6-to-7-year-old children would associate happy stickers with major melodies but not sad stickers with minor melodies (Nieminen et al., 2012). Both results are consistent with previous studies which indicated a “late” development of recognition of negative emotions in music, compared to positive emotions (Cunningham & Sterling, 1988; Dolgin & Adelson, 1990). This might be related to cognitive and intellectual maturation, as supported by the association between fluid intelligence and the ability to associate major-minor mode to happiness/sadness (Bonetti & Costa, 2019) and preference for minor-mode music (Bonetti & Costa, 2016). Conversely, a decline in cognitive and intellectual skills might underlie the reduced ability to recognize negative emotions in music in advancing age, whereas the recognition of happiness in music remains stable (Castro & Lima, 2014).

However, even adults find it difficult to identify the mode of musical excerpts in an absolute manner (Halpern et al., 1998), yet there is a significant enhancement of their performance when they are explicitly instructed on the association between major-minor mode with happiness and sadness expressions (Leaver & Halpern, 2004; Thompson & Opfer, 2014). This evidence highlights once again how mode distinction depends on both implicit and explicit learning, in addition to the individual differences in sensitivity to mode variations (Chubb et al., 2013).

#### The Role of Major-Minor Dichotomy for Emotional Regulation through Music

Music is frequently utilized for emotional regulation due to its ability to elicit diverse emotional responses (Sakka & Juslin, 2018). Given their fundamental role in shaping emotional responses to music, major and minor modes are deemed crucial in these regulation processes. Specifically, when individuals listen to music in a major key, they might experience emotions that are more uplifting and positive, leading to a sense of emotional discharge (i.e., the release or catharsis of emotions in individuals) characterized by feelings of happiness, contentment, or even euphoria. On the other hand, music in a minor key can evoke emotions that are more introspective, potentially leading to emotional discharge in the form of sadness, empathy, or a cathartic release of unexpressed negative emotions. However, these emotional responses can vary from adaptive, where music helps individuals manage and cope with emotions effectively, to maladaptive, where music exacerbates negative emotions or leads to avoidance behaviors (Carlson et al., 2015; Randall et al., 2022; Thoma et al., 2012).

Indeed, music can also lead to maladaptive emotional regulation, especially when individuals use it as a means of avoiding or suppressing negative emotions (for a review see van Goethem & Sloboda, 2011). Excessive engagement with music that reinforces negative emotions or indulges in rumination may exacerbate distress and hinder effective emotional regulation (Garrido & Schubert, 2011). In such cases, the emotional discharge experienced through music, referring to the release or catharsis of emotions in individuals, might not lead to constructive resolution but instead perpetuate negative emotional states (Taruffi & Koelsch, 2014). For instance, some individuals may find emotional discharge through minor mode music to be beneficial (Wang et al., 2018), while others may experience a deeper immersion into negative emotional states, potentially exacerbating distress (Garrido & Schubert, 2011). In this regard, it is essential to consider individual differences in emotional responses to music (e.g., Bonetti et al., 2021; Brattico et al., 2016) including potential underlying clinical conditions outlined in the next paragraph.

In conclusion, emotional regulation through music appears to depend primarily on the strategies employed by the individual (adaptive and/or maladaptive in the case of negative-valenced music), rather than the psychoacoustic characteristics of major-minor mode. However, the major-minor dichotomy provides a powerful emotional framework, allowing individuals to connect with and express various emotional states. Therefore, by selecting music in major or minor modes, individuals can tailor their emotional experience and regulation according to their emotional needs (Sachs et al., 2015).

### Clinical Observations and Implications

#### Clinical Conditions May Alter the Perception of the Major-Minor Dichotomy

Musical mode preference and perception can be significantly influenced by listener’s mood, personality profile, and psychological and/or psychiatric condition. For instance, patients with schizophrenia fail to associate minor mode music to sadness (Abe et al., 2017), as similarly seen in patients with right and left temporal resection (Khalfa et al., 2008). As regards mode preference, listeners in a sad mood do not show the typical preference for music played in major key (Hunter et al., 2011). Further, musical pieces in minor mode and slow tempo, hence with a sad connotation, tend to be more appreciated among listeners with high introversion and empathy, and low emotional stability (Ladinig & Schellenberg, 2012; Taruffi et al., 2017). In addition, high neuroticism levels are associated with stronger sad feelings in response to music and larger use of music to regulate mood and emotions (Chamorro-Premuzic et al., 2010; Ladinig & Schellenberg, 2012). Considering that neuroticism predicts both depression and anxiety disorders (Zinbarg et al., 2016), these results suggest that emotional distress might affect the preference and perception of major-minor modes.

In fact, a large body of research investigated the effect of depression on the perceived emotions in music, reporting that depression is a condition able to alter the neural responses to pleasant and favourite music (Keller et al., 2013; Osuch et al., 2009). Behavioural studies confirmed the effect of depression on emotional perception of music. Punkanen et al. (2011) found that depressed patients showed higher ratings of sadness when exposed to tender music in comparison to healthy controls, with this result highlighting a bias in which positive emotions were perceived as negative. This is in line with the results reported by Gur et al. (1992) who found that depressed patients misinterpreted neutral faces as sad and happy faces as neutral. Moreover, depressed subjects gave lower happiness ratings to happy and tender musical examples, underlying how subjects with depression slowly tend to become unresponsive to positive stimuli (Tomarkenand & Keener, 1998). A possible explanation for this tendency is the “congruency bias” which determines selective attention toward negative stimuli among depressed subjects (Gollan et al., 2008). Additionally, the effect of the negative bias appears to be correlated with the level of depression, consistent with the association between severity of depression and lower ability to discriminate happy and sad expressions suggested by Surguladze et al. (2004). Also, even a few seconds of music played in minor mode elicit emotional estimate instability (expressed as high standard deviations) in patients with retarded depression (Al’tman et al., 2000), and more intense experience in response to sad music in depressed patients compared to the healthy population was reported by Bodner et al. (2007).

Interestingly, differences in neural response to the emotional features of music occur not exclusively in major depression disorder (MDD), but also in a condition of subclinical risk of depression, which is conceptualised as an enhanced probability to develop a depressive disorder (Carlson et al., 2015). Indeed, Bonetti et al. (2017) found higher mismatch negativity (MMN) amplitude to mistuned pitches within a major context compared to MMN to pitch changes in a minor context, highlighting a different responsivity to sound frequency changes among individuals with a tendency to depression. This is consistent with the results reported by Pang et al. (2014) who recorded an impairment in the capacity to automatically process sad prosodies in depressed patients. Specifically, the MMN in response to sad syllables was absent, whereas the MMN to happy and angry prosodies was similar to the response in healthy participants.

Thus, emotional distress and depression may, in fact, impair the recognition and the processing of emotions conveyed by music, even in subclinical conditions. In this perspective, considering that major-minor dichotomy is one of the main determinants of emotional expression in music, its affective connotation might be further exploited to develop more effective and tailored music-based treatments.

### Major-Minor Dichotomy as Mood Regulator for Clinical Populations

As the major-minor dichotomy in music perception has been linked to the expression of emotional valence in music, there has also been an increasing interest in the exploration of potential clinical applications of music mode as a mood-regulating tool in disorders such as depression, anxiety, insomnia, and autism, among others.

Studies have shown that listening to music can be an effective intervention for depression, potentially improving mood and reducing symptoms. For example, a randomised controlled trial conducted by Chan et al. (2012) found that listening to music resulted in significant reductions in depressive symptoms among patients with major depressive disorder. Moreover, a systematic review by Maratos et al. (2008) suggested that listening to self-selected music was effective in reducing symptoms of depression and anxiety in a group of older adults. In addition to passive listening, active engagement with music in a minor key can also be a therapeutic approach for depression. For example, a study by Erkkilä et al. (2011) found that improvising on a keyboard was effective in reducing symptoms of depression and anxiety among psychiatric patients.

Thus, the use of musical mode as an intervention for depression is thought to be related to the emotional and psychological responses that different modes can elicit. For example, minor keys may be particularly effective for individuals who have difficulty expressing or processing negative emotions, as they can provide a safe and non-threatening way to engage with these emotions, leading to a sense of emotional release and relief. Indeed, minor music has been linked to a better psychological regulation and relaxation compared to major music Wang et al. (2018). Additionally, the dichotomy may be helpful for individuals who experience symptoms such as rumination or excessive worry, as they can provide a way to shift focus away from negative thoughts and emotions (Garrido et al., 2017).

Furthermore, research has shown that music may be effective in reducing arousal levels in individuals with insomnia. There are various factors that can contribute to insomnia, including stress, anxiety, and depression. One theory that explains the mechanism behind insomnia is the hyperarousal theory, which suggests that individuals with insomnia have a heightened level of arousal that prevents them from falling asleep or staying asleep. A meta-analysis of randomised controlled trials by Jespersen et al. (2022) showed moderate-certainty evidence that listening to music improves sleep quality when compared to no treatment. Therefore, music in the minor mode may also be used to regulate and ameliorate the state of hyperarousal in individuals with insomnia, however, this hypothesis remains untested.

Moreover, the use of the mode dichotomy may be beneficial for children with autism spectrum disorder (ASD) who typically show altered processing of affective information within social and interpersonal domains. In fact, a study by Heaton et al. (1999) showed that children with ASD and age and intelligence matched controls did not differ in their ability to ascribe affective connotations (happy/sad faces) to melodies in the major or minor mode. Therefore, the dichotomy could have potential benefits as music therapy for ASD by helping them to learn, recognize, and differentiate between different emotional states, regulate their own emotions, communicate effectively, integrate, and regulate sensory information, and develop social skills.

Overall, the major-minor dichotomy in music perception is a significant determinant of emotional expression in music and is influenced by individual factors such as mood, personality, and clinical conditions.

Emotional distress and depression can impair the recognition and processing of emotions conveyed by music, including the major-minor dichotomy. However, the use of musical mode as an intervention for depression and other clinical conditions has shown promise, with minor keys being particularly effective for individuals who have difficulty expressing or processing negative emotions. Overall, further research is needed to better understand the mechanisms behind the relationship between music and emotional regulation in clinical populations. Nevertheless, the potential clinical applications of music mode highlight the importance of music in promoting emotional well-being and suggest its potential as a tool for supporting mental health.

### The Major-Minor Mode(l)

Music has been an important part of human life for centuries, and it has been linked to emotional expression and regulation. In particular, the major-minor dichotomy has been shown to be a powerful tool in the perception and expression of emotional valence in music. This dichotomy is influenced by various individual factors, such as age, cultural background, musical expertise, mood, personality, and clinical conditions such as depression.

Based on the findings of this systematic review, we propose a comprehensive model of musical mode perception and its behavioural and physiological correlates that incorporates these factors and their interactions to better understand the complex emotional responses that major and minor musical modes elicit (**Figure 5**).

**Figure 5.**
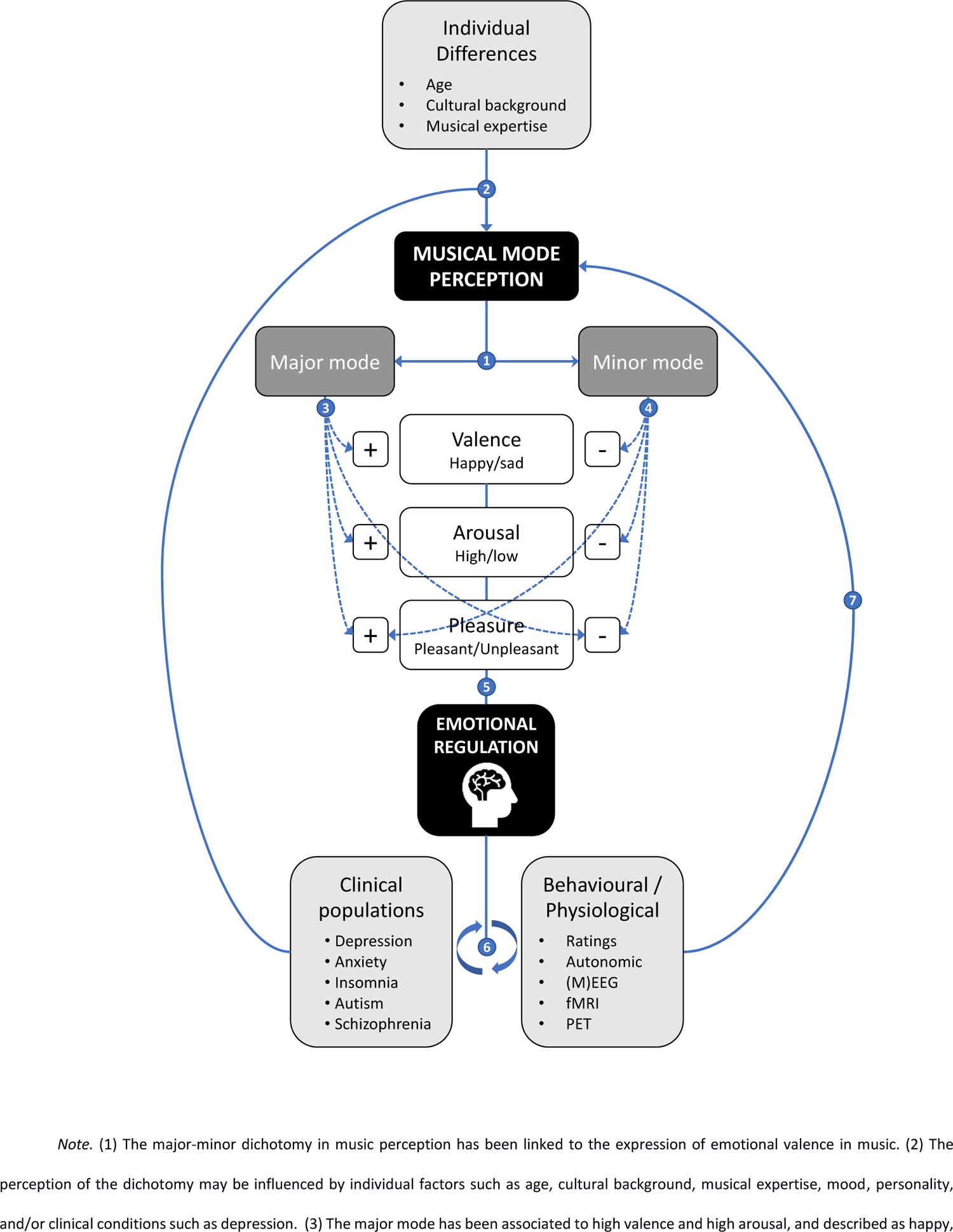
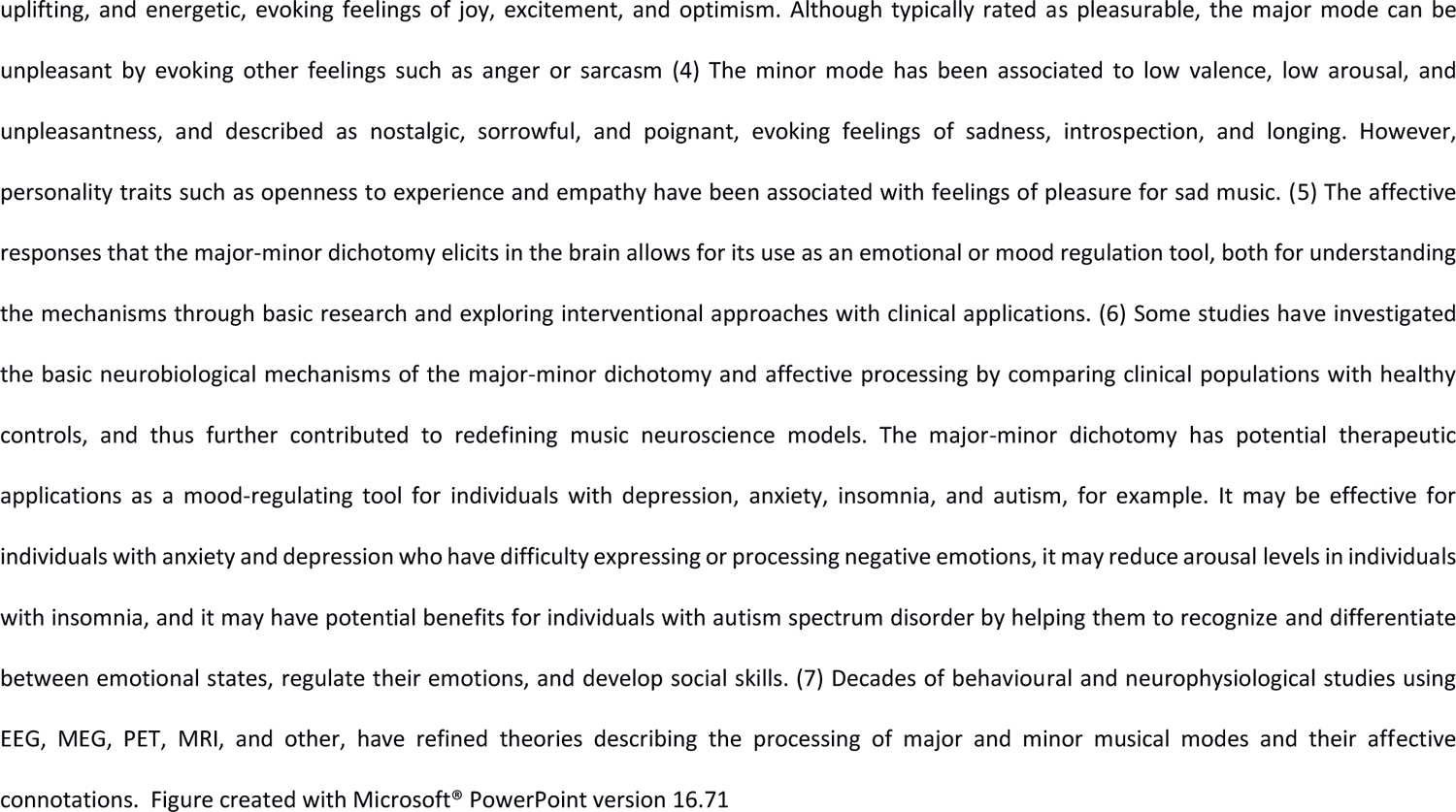
The Major-Minor Model(I)

### The Major-Minor Dichotomy

The major-minor dichotomy is a fundamental aspect of Western tonal music, which divides musical modes into two categories: major and minor. The major mode is often associated with positive emotions, such as happiness, excitement, and optimism, while the minor mode is associated with negative emotions, such as sadness, introspection, and longing. However, it is important to note that major and minor modes can evoke a wide range of emotions, and they are not limited to the basic affective connotations. The major-minor dichotomy is a complex and multifaceted phenomenon, and it is influenced by various individual and cultural factors.

### Individual Factors

Individual factors such as age, cultural background, musical expertise, mood, personality, and clinical conditions such as depression can influence the perception of the major-minor dichotomy in music. Personality traits such as empathy and openness to experience may also affect how individuals respond to sad music in the minor mode. Cultural factors may influence the perceived emotional valence of different musical modes.

### Behavioural and Brain Correlates

Over the course of several decades, multiple neuroimaging techniques including EEG, MEG, PET, and MRI have been used in numerous studies to improve our understanding of the cognitive and neural processes involved in perceiving major and minor modes in music. Additionally, comparisons between clinical groups and healthy individuals have also played a role in refining music neuroscience models.

### Clinical Implications

The major-minor dichotomy in music perception has been explored as a potential mood-regulating tool for clinical populations, including those with depression, anxiety, insomnia, and autism. Listening to music has been shown to be an effective intervention for depression, with minor keys providing a safe way to engage with negative emotions. Major-minor dichotomy can also be exploited to reduce arousal levels in individuals with insomnia and has potential as a music therapy for children with ASD in terms of emotion recognition and social skills. Further research is needed to understand the relationship between music and emotional regulation in clinical populations, but the potential applications highlight the importance of music in promoting emotional well-being.

### Limitations and Future Directions

Published studies exploring the perception of major-minor dichotomy are several and difficult to synthesise due to differences in methodologies, stimuli, and population. This probably underlies the discrepancies in the pattern of results among the papers we reviewed, especially concerning neuroimaging studies. From this work, we offer some recommendations for future directions that empirical research on major and minor modes could pursue, and we discuss the potential implications.

Primarily, we highlighted how intrapersonal factors may substantially contribute to the recognition and emotional connotation attributed to major and minor modes. Therefore, we suggest that future studies aiming at exploring the neural and physiological response to music mode should always consider the subjective psychological perception of pleasantness and listener’s level of musical expertise, since these factors may significantly affect brain activation and perceived emotions in response to stimuli differing in mode. The combination of subjective questionnaires and physiological measures would provide reliable information about the effective consonance between physiological state and perceived emotion.

On this note, we also suggest that future studies in musical mode and music in general, should include more information about the participants. We found that very few of the included studies reported years of education, year of musical training, onset of musical training, and current music practice time. Furthermore, we found, through the quality assessment, that items related to subject characterization and confounding effects got lower scores. Including such variables in scientific studies of human behaviour and music neuroscience allows researchers to control for individual differences, measure expertise, investigate developmental effects, and understand plasticity in response to musical training (Criscuolo et al., 2022).

A further crucial question in musical mode investigation concerns the type of stimulation to employ. For instance, some studies here reviewed used musical chords as stimuli, namely single sounds, whereas other studies, to better capture the complex nature of music, exposed participants to longer musical excerpts or even whole songs in major or minor keys. In the latter case, it appears quite complicated to identify the specific role of musical mode in determining the emotional response, since other variables such as tempo and lyrics do affect the perceived emotions in music (Brattico et al., 2011; Peretz et al., 1998). Also, during the exposure to longer musical stimuli, motivation and attention are more likely to influence the subjective responses. Accordingly, papers employing chords or short excerpts showed higher consistency of results than those eliciting responses with longer or less natural harmonic sounds. However, longer real-life musical stimuli seem to be more appropriate than shorter sounds when the investigation is focused on the experienced mood (McConnell & Shore, 2011), or on the association with personality traits (Dobrota & Reić Ercegovac, 2015; Ladinig & Schellenberg, 2012). In fact, longer sounds are necessary to induce and maintain an emotional response in the listener (Eerola & Vuoskoski, 2013), as well as to allow for an evaluation of elicited responses overtime during the exposure. On the contrary, in addition to help reducing the effect of other musical features, shorter stimuli are more suitable when the investigation is aimed at evaluating basic emotions, since as reported by Peretz (2001) a very short duration of music (250 ms) is enough to differentiate sadness from happiness. Therefore, exploring the emotional response to major-minor music in terms of the happiness-sadness dichotomy seems justified when stimuli are brief (such as musical chords). On the other hand, studies using real musical pieces, hence involving multiple elements and structural complexities that can influence emotional experiences, should consider more nuanced and complex emotional responses, such as wonder, sublimity, awe, and nostalgia. Indeed, only few studies included in our review assessed beyond happy-sad/pleasant-unpleasant participants’ responses, revealing in this regard a gap in research. Along this line, further explorations might also directly compare emotional responses elicited by single chords to those induced by longer musical pieces to understand how the length of stimuli affects emotional perception. In general, to identify the effect of major-minor dichotomy in a context of either longer or shorter stimuli, it is suggested to minimise the variations of other musical features during the exposure, limiting the changes solely to musical mode.

When assessing publication bias in our meta-analyses, we found that the funnel plots for both behavioural and EEG meta-analyses showed a symmetric distribution. For ALE meta-analyses, using the Fail Safe-N analysis, we found adequate robustness of our results, with only one ROI showing an FSN below the minimum imposed in each of the ALE within contrasts (CAU-R), thus, indicating an overall robust convergence of brain areas in our study (**Table S7**).

Lastly, the findings of this study have several implications for future research on musical mode perception also in a clinical perspective. For instance, this systematic review reveals that the preference towards music in minor mode is often aimed at regulating mood and providing emotional benefits, but it might also underlie a condition of emotional distress which, in turn can lead to pathological conditions (Garrido & Schubert, 2011; Ladinig & Schellenberg, 2012; Taruffi & Koelsch, 2014). Consistently, prior evidence indicated that depression may affect the emotional evaluation of musical stimuli by eliciting lower responsivity to positive stimuli as well as eliciting a stronger response to negative stimuli (Bodner et al., 2007; Punkanen et al., 2011).

Specifically, the preference for minor music might serve as an early indicator of emotional distress, such as in case of depressed individuals. Similar results might also be obtained associating the performance of emotional recognition of musical mode with the listeners’ level of anxiety, since previous studies suggested that also patients with anxiety show a negative bias in emotion recognition tasks (Quadflieg et al., 2007). Given the above, the affective evaluation of musical mode fragments might help to identify emotional disorders, with the advantages of (1) not involving complex cognitive mediation, since it implies a rapid evaluation of short sounds, (2) bypassing the verbal barriers in assessment and (3), exploiting the generally positive attitude towards exposure to music. Moreover, if on one side there is evidence of large and systematic effect of individual differences in major-minor distinction, which might make it difficult to use judgments of mode to identify emotional disorders, on the other, studies employing shorter and unambiguous stimuli, such as diatonic scales (Tan & Temperley, 2017) or triad chords (Bakker & Martin, 2015; Pallesen et al., 2003) seem to limit this tendency. In this perspective, the use of musical mode as a clinical stimulation paradigm is worth further investigation. Furthermore, neuroimaging evidence of the effect of depression level in the automatic discrimination of major-minor stimuli, as indexed by the auditory component of mismatch negativity (MMN) (Bonetti et al., 2017) suggests that the preference for sad music among depressed subjects could have a basis or, at least, a neural correlate in the sensory processes on the auditory cortex level.

## Conclusions

The major-minor dichotomy has long been of interest in music research studies. Although extensive literature exists exploring this fundamental musical feature, this is the first attempt to comprehensively and systematically review and synthesise research specifically investigating physiological and behavioural correlates of major and minor modes, as well as summarise how this dichotomy has been investigated in empirical research. The papers were analysed both in qualitative terms and through three different quantitative meta-analysis, based on the methods applied in the studies. The quantitative synthesis allowed us to point out the distinct behavioural and neural responses elicited by major and minor mode in music.

The typical affective connotation attributed to the major-minor dichotomy, hence major-happy and minor-sad association, has been substantially confirmed in the examined literature, including cross-cultural studies which hence suggest a universal component of this affective categorization (Fang et al., 2017; Hoshino, 1996; Mao & Rau, 2014). In other words, the emotional dimension associated with major and minor modes might primarily arise from their different psychoacoustic features, such as harmonicity, spectral entropy, and roughness, rather than from cultural factors. In line with this view, some authors speculated that minor mode music chords share similar harmonic relationships to sad speech prosody. Nonetheless, empirical research aimed at establishing the universality of major/minor affective dimension is increasingly hindered by the difficulty in finding population groups totally unfamiliar with Western culture and music (Virtala & Tervaniemi, 2017). In the latter case, the typical affective perception of major and minor appears significantly reduced (Fang et al., 2017; Hoshino, 1996). A comprehensive view might be that psychoacoustics characteristics predispose to the development of the conventional affective association of major with happiness, and minor with sadness, leading to established cultural-based emotional connotations. Subsequently, levels, and types of exposure to Western-like music might contribute to these associations by modulating familiarity and the ensuing ability to discriminate between the two modes.

Importantly, the perception of distinct emotions in music (such as happiness-sadness) does not always correspond to the subjective emotional response (such as liking, pleasantness). This distinction is particularly relevant, especially for the minor mode, as despite its strong negative connotation, it often elicits pleasant emotions and even positive implications in terms of emotional regulation. In this regard, studies specifically addressing major-minor dichotomy in music has not always considered the “aesthetic” potential of minor music, limiting exploration to its strictly negative connotation. However, in music, it is not uncommon for the negative connotation to be used as a sort of springboard to achieve a more impactful emotional effect. For instance, in this review we indicated some famous musical pieces employ transitioning from minor to major mode to accentuate a joyful and triumphant conclusion.

The centrality of the subjective emotional response to major-minor processing is confirmed by neuroimaging studies which, indeed, point out a crucial point: neural and physiological activity appears to be more associated with basic emotional responses and evaluation of pleasantness rather than musical mode itself (Lee et al., 2011; Mizuno & Sugishita, 2007; Suzuki et al., 2008). This may partly underlie the discordant results that arose in studies investigating the neural correlates of mode perception, in addition to the differences of methods and stimuli. Again, the reviewed studies seem to emphasize a substantial difference between the emotional expression of music and the emotional responses evoked.

In this perspective, any study exploring the realm of major-minor dichotomy cannot disregard the investigation of the intra-personal factors contributing to its perception and processing, as pointed out in our proposed model. For instance, in this review we reported several studies suggesting that individual difference in processing of major-minor mode partly arises from musical expertise, since musicians show more neural activity and appreciation for minor-mode music than non-musicians (Brattico et al., 2016; Halpern et al., 2008; Khalfa et al., 2005) as well as a greater ability to discriminate between the two modes (Halpern et al., 2008; Hoshino, 1996; Leaver & Halpern, 2004). Therefore, studies reviewed agree in considering musical expertise a determinant of the major-minor representation and its affective processing as well as a factor which significantly speeds up the musical mode categorization process in children (Virtala et al., 2012). The fact that major-minor perception is mostly related to its affective connotation might also explain why its explicit perception and affective recognition (mainly regarding minor/sad association) are established only after the age of 3-4, concomitantly with cognitive and intellectual maturation, such as fluid intelligence (Bonetti & Costa, 2019; Dalla Bella et al., 2001; Kastner & Crowder, 1990). Accordingly, ageing leads to a reduced ability to perceive the emotional connotation of musical mode, especially concerning negative emotions conveyed by minor mode excerpts (Dalla Bella et al., 2001). In other words, the ability to explicitly perceive (and supposedly discriminate) major and minor modes is strongly related to the ability of recognizing their emotional connotation, supporting the conjecture that the ensuing behavioural and physiological responses are not related to the musical mode *per se*.

In conclusion, this work has aimed to provide a comprehensive and in-depth understanding of one of the main features of Western music, characterized by a prominent and dichotomous affective connotation. One of the main challenges in studying a specific musical feature such as the major-minor dichotomy lies in the difficulty of assessing its affective impact without considering all the other elements inherent to music, such as melody, harmony, tempo, and dynamics, in addition to individual differences. In music, all these features interact with each other to evoke specific and unique emotional responses (Juslin & Sloboda, 2013). In this sense, music aligns to the popular saying that *the whole is greater than the sum of its parts*, as various features combine to create a more meaningful and powerful emotional experience. Therefore, while exploring the individual components that constitute music is crucial for grasping its underlying principles, it may fall short in capturing the intricate and profound emotional experience that music elicits.

Nevertheless, musical mode has been a central topic in musical studies since Greek theorists, including Gregorian, Renaissance-polyphony and tonal-harmonic studies (Powers et al., 2001). The phenomenon that mainly captured the attention of theorists and musicians was how the structure of musical scales related to their specific emotional expression, a phenomenon that was largely demonstrated in controlled studies, as reported in this systematic review. Although the literature on musical mode has reached a voluminous corpus that we have summarised in this review, we still miss a comprehensive explanation of the reasons why different modes are associated with different emotional expressions in music. The study of musical mode, in this perspective, could be central for gaining significant insights into the central theme of emotional valence perception and processing in affect theory.

