## Supplementary material for "The Major-Minor mode Dichotomy in Music Perception: A Systematic Review and Meta-Analysis on its Behavioural, Physiological, and Clinical Correlates"

### Table of Contents

**Search strings**[1] 2

**Figure S1.** ***PRISMA Flowchart for Literature Search Process***[2] 3

**Table S1.** ***PRISMA Checklist***[2] 4

**Table S2.** ***Main Results of Included Studies*** 6

**Table S3.** ***Quality Assessment of All included Studies***[3] 12

**Table S4.** ***fMRI Acquisition and Analysis*** 15

**Table S5.** ***Functional Characterization of ROIs as Included on BrainMap Platform Resulted from ALE Meta-analyses*** 16

**Table S6.** ***Meta-analytic Connectivity Modelling of ROIs Resulted from ALE Meta-analyses.*** 18

**Table S7.** ***FSN Robustness Assessment of ROIs Resulted from ALE Meta-analyses*** 20

#### Search strings[1]

We included two separate searches and screenings, after recommendations by a previous peer-review process.

Previous search: May 2018
(musical mode OR musical modes OR music mode OR music modes) AND (perception OR psychology OR behavior OR behaviour OR brain OR cerebral OR emotion OR emotions OR emotional OR happiness OR sadness OR depression OR cognition OR auditory perception OR chords OR electroencephalography OR neural correlates OR neural correlate OR mismatch negativity OR cortex OR cerebral cortex OR magnetoencephalography OR magnetic resonance imaging OR children OR child OR infant OR infants OR adult OR adults OR limbic system)

Current last search: August 2023

**PubMed**
("music"[MeSH Terms] OR "musician"[All Fields] OR "music"[All Fields] OR "music listening" [All Fields] OR "music production" [All Fields] OR "music performance" [All Fields] OR "music playing" [All Fields] OR "music composition" [All Fields] OR "music perception" [All Fields] OR "musical mode"[All Fields] OR "major mode"[All Fields] OR "minor mode"[All Fields] OR "musical training"[All Fields] OR "musical expertise"[All Fields]) AND ("emotions"[MeSH Terms] OR "valence"[All Fields] OR "arousal"[MeSH Terms] OR "happiness"[MeSH Terms] OR "sadness"[MeSH Terms] OR "mood disorders"[MeSH Terms] OR "depression"[MeSH Terms]) AND ("magnetic resonance imaging"[MeSH Terms] OR "electroencephalography"[MeSH Terms] OR "evoked potentials"[MeSH Terms] OR "magnetoencephalography"[MeSH Terms] OR "transcranial magnetic stimulation"[MeSH Terms] OR "electrocorticography "[MeSH Terms])
614

**Scopus**
(TITLE-ABS-KEY("music") OR TITLE-ABS-KEY("musician") OR TITLE-ABS-KEY("music listening") OR TITLE-ABS-KEY("music production") OR TITLE-ABS-KEY("music performance") OR TITLE-ABS-KEY("music playing") OR TITLE-ABS-KEY("music composition") OR TITLE-ABS-KEY("music perception") OR TITLE-ABS-KEY("musical mode") OR TITLE-ABS-KEY("major mode") OR TITLE-ABS-KEY("minor mode") OR TITLE-ABS-KEY("musical training") OR TITLE-ABS-KEY("musical expertise")) AND (TITLE-ABS-KEY("emotions") OR TITLE-ABS-KEY("valence") OR TITLE-ABS-KEY("arousal") OR TITLE-ABS-KEY("mood disorders") OR TITLE-ABS-KEY("depression")) AND (TITLE-ABS-KEY("magnetic resonance imaging") OR TITLE-ABS-KEY("electroencephalography") OR TITLE-ABS-KEY("evoked potentials") OR TITLE-ABS-KEY("magnetoencephalography") OR TITLE-ABS-KEY("transcranial magnetic stimulation") OR TITLE-ABS-KEY("electrocorticography"))
882

**PsycInfo**
(Music OR Musician OR Music listening OR Music production OR Music performance OR Music playing OR Music composition OR Music perception OR Musical mode OR Major mode OR Minor mode OR musical training OR Musical expertise) AND (Emotions OR Valence OR Arousal OR Mood disorders OR Depression) AND (Magnetic resonance imaging OR Electroencephalography OR Evoked potentials OR Magnetoencephalography OR Transcranial magnetic stimulation OR Electrocorticography)
875

#### Figure S1

*PRISMA Flowchart for Literature Search Process*[2]

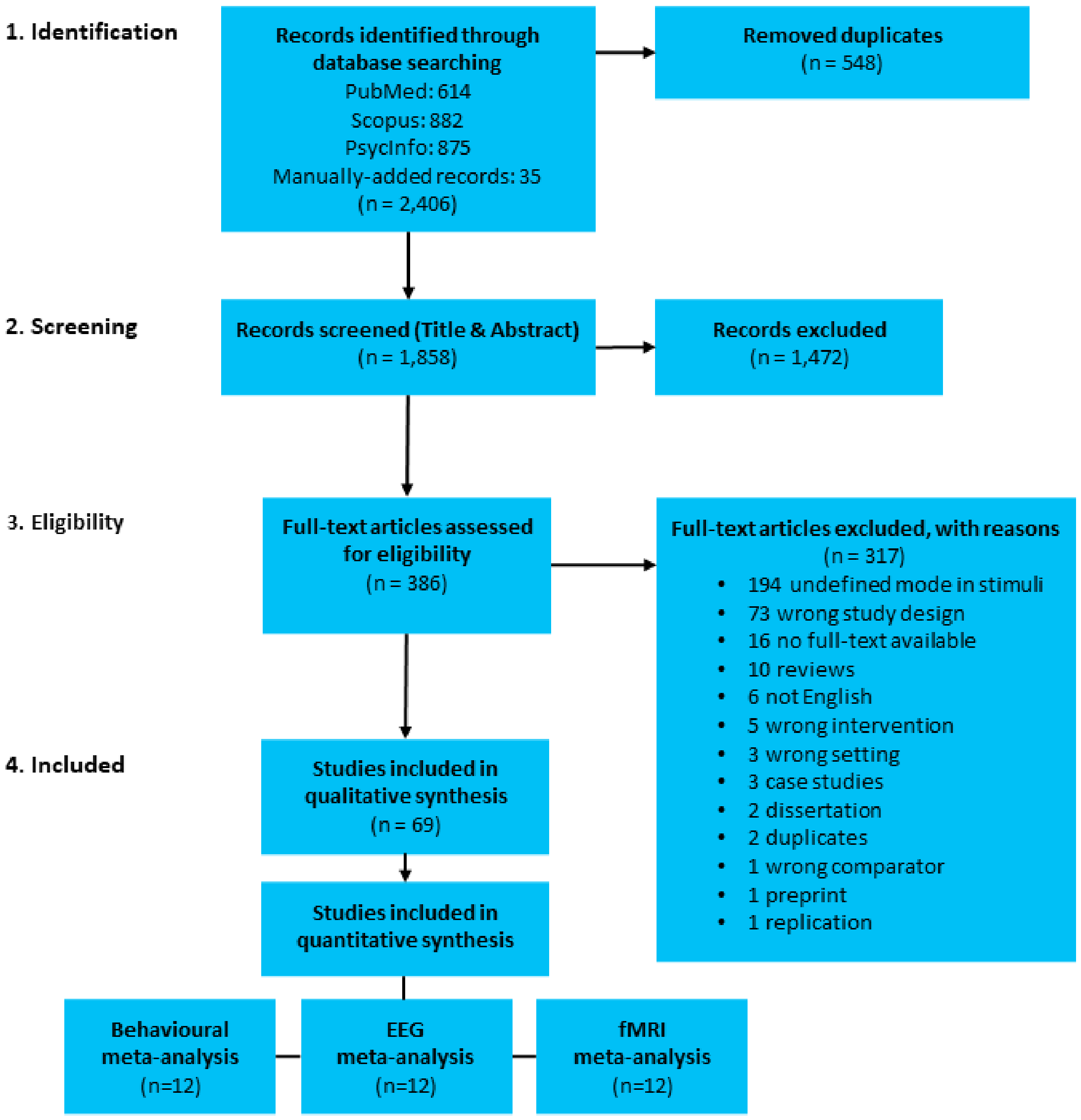


**Supplementary Figure 1**. PRISMA flowchart for literature search process.

**Table S1**

*PRISMA Checklist*[2]

SI = Supplementary Information

| Section/topic | # | Checklist item | Reported on page # |
|---|---|---|---|
| **TITLE** | | | |
| Title | 1 | Identify the report as a systematic review, meta-analysis, or both. | 1 |
| **ABSTRACT** | | | |
| Structured summary | 2 | Provide a structured summary including, as applicable: background; objectives; data sources; study eligibility criteria, participants, and interventions; study appraisal and synthesis methods; results; limitations; conclusions and implications of key findings; systematic review registration number. | 2 |
| **INTRODUCTION** | | | |
| Rationale | 3 | Describe the rationale for the review in the context of what is already known. | 4 |
| Objectives | 4 | Provide an explicit statement of questions being addressed with reference to participants, interventions, comparisons, outcomes, and study design (PICOS). | 5 |
| **METHODS** | | | |
| Protocol and registration | 5 | Indicate if a review protocol exists, if and where it can be accessed (e.g., Web address), and, if available, provide registration information including registration number. | 6 |
| Eligibility criteria | 6 | Specify study characteristics (e.g., PICOS, length of follow-up) and report characteristics (e.g., years considered, language, publication status) used as criteria for eligibility, giving rationale. | 7 |
| Information sources | 7 | Describe all information sources (e.g., databases with dates of coverage, contact with study authors to identify additional studies) in the search and date last searched. | 7 |
| Search | 8 | Present full electronic search strategy for at least one database, including any limits used, such that it could be repeated. | 7 |
| Study selection | 9 | State the process for selecting studies (i.e., screening, eligibility, included in systematic review, and, if applicable, included in the meta-analysis). | 7-8 |
| Data collection process | 10 | Describe method of data extraction from reports (e.g., piloted forms, independently, in duplicate) and any processes for obtaining and confirming data from investigators. | 8-9 |
| Data items | 11 | List and define all variables for which data were sought (e.g., PICOS, funding sources) and any assumptions and simplifications made. | 8-9 |
| Risk of bias in individual studies | 12 | Describe methods used for assessing risk of bias of individual studies (including specification of whether this was done at the study or outcome level), and how this information is to be used in any data synthesis. | 8 |
| Summary measures | 13 | State the principal summary measures (e.g., risk ratio, difference in means). | NA |
| Synthesis of results | 14 | Describe the methods of handling data and combining results of studies, if done, including measures of consistency (e.g., $I^2$) for each meta-analysis. | 9-10-11 |

| Section/topic | # | Checklist item | Reported on page # |
|---|---|---|---|
| Risk of bias across studies | 15 | Specify any assessment of risk of bias that may affect the cumulative evidence (e.g., publication bias, selective reporting within studies). | 8 |
| Additional analyses | 16 | Describe methods of additional analyses (e.g., sensitivity or subgroup analyses, meta-regression), if done, indicating which were pre-specified. | NA |
| **RESULTS** | | | |
| Study selection | 17 | Give numbers of studies screened, assessed for eligibility, and included in the review, with reasons for exclusions at each stage, ideally with a flow diagram. | 13 |
| Study characteristics | 18 | For each study, present characteristics for which data were extracted (e.g., study size, PICOS, follow-up period) and provide the citations. | 13 |
| Risk of bias within studies | 19 | Present data on risk of bias of each study and, if available, any outcome level assessment (see item 12). | 15 |
| Results of individual studies | 20 | For all outcomes considered (benefits or harms), present, for each study: (a) simple summary data for each intervention group (b) effect estimates and confidence intervals, ideally with a forest plot. | 13 |
| Synthesis of results | 21 | Present results of each meta-analysis done, including confidence intervals and measures of consistency. | 15-16-17-18 |
| Risk of bias across studies | 22 | Present results of any assessment of risk of bias across studies (see Item 15). | 15 |
| Additional analysis | 23 | Give results of additional analyses, if done (e.g., sensitivity or subgroup analyses, meta-regression [see Item 16]). | NA |
| **DISCUSSION** | | | |
| Summary of evidence | 24 | Summarize the main findings including the strength of evidence for each main outcome; consider their relevance to key groups (e.g., healthcare providers, users, and policy makers). | 19 |
| Limitations | 25 | Discuss limitations at study and outcome level (e.g., risk of bias), and at review-level (e.g., incomplete retrieval of identified research, reporting bias). | 42 |
| Conclusions | 26 | Provide a general interpretation of the results in the context of other evidence, and implications for future research. | 46 |
| **FUNDING** | | | |
| Funding | 27 | Describe sources of funding for the systematic review and other support (e.g., supply of data); role of funders for the systematic review. | NA |

**Table S2**

*Main Results of Included Studies*

| | Author (year) | Title | Main results |
|---|---|---|---|
| 1 | *Abe et al. (2017)* | Music-evoked emotions in schizophrenia | Patients diagnosed with schizophrenia significantly failed to recognize minor mode chords as sad compared to healthy controls. |
| 2 | *Al'tmann et al. (2000)* | Estimation of short musical fragments in normal subjects and patients with chronic depression | Compositions in a minor key determined emotional estimate instability (expressed as high standard deviations) in patients with retarded depression. |
| 3 | *Bakker & Martin (2015)* | Musical chords and emotion: Major and minor triads are processed for emotion | N2 amplitudes were significantly lower for congruent (major mode–happy faces and minor mode–sad faces) than for incongruent pairings. No significant effects of chord congruence were observed for the P1. |
| 4 | *Bonetti & Costa (2016)* | Intelligence and musical mode preference | Preference for minor mode is significantly associated with fluid intelligence. |
| 5 | *Bonetti & Costa (2019)* | Musical mode and visual- spatial cross-modal associations in infants and adult | Children develop the association minor mode/sadness later than major mode/happiness. Warm-cold colours were associated with happy-sad stimuli in adults but not in children. |
| 6 | *Bonetti et al. (2017)* | Risk of depression enhances auditory Pitch discrimination in the brain as indexed by the mismatch negativity | Results showed higher MMN amplitudes to mistuned pitches within a major context compared to MMN to pitch changes in a minor context, suggesting a higher responsivity to sound frequency changes also in a subclinical condition of depression. |
| 7 | *Brattico et al. (2009)* | Neural discrimination of nonprototypical chords in music experts and laymen: An MEG study | MMN measures indicated an enhanced reactivity of the auditory cortex to unpleasant chords in musicians, but no difference was found in MMN to minor chords between musicians and non-musicians. |
| 8 | *Brattico et al. (2011)* | A functional MRI study of happy and sad emotions in music with and without lyrics | Sad music (in minor mode) induced activity within the right caudate head and the left thalamus, whereas instrumental happy music (in major mode) significantly activated structures of the limbic system and the right pars opercularis of the inferior frontal gyrus, with the involvement of auditory regions only to happy music with lyrics. |
| 9 | *Brattico et al. (2016)* | It's sad but I like it: The neural dissociation between musical emotions and liking in experts and laypersons | Well-trained musicians consider music in minor mode as beautiful and pleasant compared to non-musicians. |
| 10 | *Castro & Lima (2014)* | Age and musical expertise influence emotion recognition in music | Music training is associated with a more accurate categorization of musical emotions in response to sad/happy musical excerpts. |

| | | | |
|---|---|---|---|
| 11 | *Chang et al. (2015)* | Experiencing affective music in eyes-closed and eyes-open states: an electroencephalography study | When listening to musical pieces with different emotional content (i.e., positive music in major mode and negative music in minor mode), participants in eyes-open condition tend to rate music as more positive-valenced compared to eyes-closed condition. Also, the theta power in the frontal area significantly increased while listening to emotional-positive music compared to emotional-negative music under the eyes-closed condition. |
| 12 | *Chubb et al. (2013)* | Bimodal distribution of performance in discriminating major/minor modes | When asked to classify single tone-scrambles as major or minor, approximately 70% of listeners performed near chance, whereas the other 30% performed nearly perfectly. |
| 13 | *Costa (2013)* | Effects of mode, consonance, and register in visual and word-evaluation affective priming experiments | Mode can automatically influence non-musical cognitive tasks such as word and picture evaluation. When presented with an effectively congruent chord as prime, target words or pictures are evaluated faster and more correctly. |
| 14 | *Costa et al. (2004)* | Interval distributions, mode, and tonal strength of melodies as predictors of perceived emotion | The attribution of happiness and serenity is associated with the major mode. |
| 15 | *Dalla Bella et al. (2001)* | A developmental study of the affective value of tempo and mode in music | 5-year-olds children can discriminate happy and sad musical excerpts, but they only used information about tempo, whereas starting from 6-8 years old they utilized both tempo and mode, like adults. |
| 16 | *Dobrota & Ervegovac (2015)* | The relationship between music preferences of different mode and tempo and personality traits-implications for music pedagogy | Emotional stability and optimism are significant predictors of preferences towards music in major mode. |
| 17 | *Eerola et al. (2013)* | Emotional expression in music: Contribution, linearity, and additivity of primary musical cues | When listeners judged the perceived emotional characters of 200 musical examples, mode was the most important cue. |
| 18 | *Ellison et al. (2015)* | Affective versus cognitive responses to musical chords: An ERP and behavioral study | In an ERP study, participants classified major and minor chords as happy and sad, respectively. Moreover, sad-rated, and minor chords were followed by negative ERPs at 300 ms postonset. |
| 19 | *Fang et al. (2017)* | Perception of Western musical modes: A Chinese study | Results show that major mode music among Chinese people induces greater pleasure, arousal, and liking ratings than minor-mode music, which instead induces greater tension. |
| 20 | *Franco et al. (2017)* | Preschoolers' attribution of affect to music: A comparison between vocal and instrumental performance | The results showed that even the youngest children were able to correctly identify the intended emotion in music, though the recognition of happy music appeared facilitated in the instrumental condition. |
| 21 | *Gosselin et al. (2005)* | Impaired recognition of scary music following unilateral temporal lobe excision | Recognition of happy (major mode) and sad (minor mode) musical excerpts was not impaired in patients with unilateral medial temporal lobe excision. |
| 22 | *Green et al (2008)* | Music in minor activates limbic structures: a relationship with dissonance? | Minor melodies caused increased activity in limbic structures, namely left parahippocampal gyrus, bilateral ventral anterior cingulate, and in left medial prefrontal cortex compared to major melodies. Minor mode music was rated as sadder than major mode, whereas no significant difference was found in liking ratings. |
| 23 | *Gregory et al. (1996)* | The development of emotional responses to music in young children | 7 to 8-year-olds children (like adults) showed significant differential responses to major and minor mode, but 3 to 4- year-olds children showed almost no difference, suggesting that the emotional response related to musical mode develops between the ages of 4 and 7 years. |

| | | | |
|---|---|---|---|
| *24* | *Halpern (1984)* | Perception of structure in novel music | When asked to discriminate between tunes in major and minor mode, non-musicians perform worse than musicians. |
| *25* | *Halpern et al. (1998)* | Perception of mode, rhythm, and contour in unfamiliar melodies: Effects of age and experience. | The discrimination of major from minor tunes is difficult for everyone, even for musicians. |
| *26* | *Halpern et al. (2008)* | An ERP study of major- minor classification in melodies | When asked to judge each melody as major/minor and happy/sad, musicians showed a late P3 to the critical note only for minor melodies in both tasks. Moreover, non-musicians who did not have affective labels available were unable to do this classification, and non-musicians who either were told about affective labels, showed moderately good performance. |
| *27* | *Heaton et al. (1999)* | Can children with autistic spectrum disorders perceive affect in music? An experimental investigation | No significant difference arose between children with autism and matched controls in identifying the affective connotations of major and minor mode music. |
| *28* | *Hevner (1935)* | The affective character of the major and minor modes in music | Major mode was associated with "happy" adjectives whilst the participants selected more "sad" adjectives when the music was played in minor mode. |
| *29* | *Hoshino (1996)* | The feeling of musical mode and its emotional character in a melody | Even non-western listeners associate the major mode with positive emotions and the minor mode with negative emotions. |
| *30* | *Hunter et al. (2008)* | Mixed affective responses to music with conflicting cues | Listeners tend to find music they rate as happy (major key) also as pleasant, whereas music rated as sad is perceived as unpleasant. |
| *31* | *Hunter et al. (2011)* | Misery loves company: Mood-congruent emotional responding to music | After inducing sad mood, the typical preference for major mode reduced and ambiguous stimuli are perceived as sadder. |
| *32* | *Jenni et al. (2017)* | Impact of major and minor mode on EEG frequency range activities of music processing as a function of expertise | Minor and major compositions distinctively modulated synchronisation of neuronal activities in high-frequency ranges (beta and gamma) in frontal regions, with increased activity in response to minor compositions especially in musicians and in experts. High-frequency electroencephalographic (EEG) activities carry information about musical mode. |
| *33* | *Jeong et al. (2011)* | Congruence of happy and sad emotion in music and faces modifies cortical audiovisual activation | The activation of superior temporal gyrus (STG) when happy music (major mode) was presented with happy faces was greater than the activation seen when sad music (minor mode) was presented with sad faces. In contrast, incongruent stimuli diminished the BOLD response in STG and elicited greater signal change in bilateral FG. When presented with happy music, happy faces were rated as happier, and sad faces were rated as less sad. When presented with sad music, happy faces were rated as less happy, and sad faces were rated as sadder. |
| *34* | *Jomori et al. (2013* | Effects of emotional music on visual processes in inferior temporal area | A relatively larger M170 response during visual stimulation in the inferior temporal area was induced during negative (minor mode) compared to positive (major mode) music. |
| *35* | *Kastner & Crowder (1990)* | Perception of the major/minor distinction: IV. Emotional connotations in young children | Children aged 3 and 12 years, showed both a reliable positive/major negative/minor association. |
| *36* | *Khalfa et al. (2005)* | Brain regions involved in the recognition of happiness and sadness in music | Results of the minor versus major mode contrast showed a larger activity in the posterior cingulate cortex, and in the left orbital and mid-dorsolateral frontal cortex. |

| | | | |
|---|---|---|---|
| 37 | *Khalfa et al. (2008)* | Evidence of lateralized anteromedial temporal structures involvement in musical emotion processing | Both right and left temporal resections impaired sadness (minor mode music) recognition, whereas happiness (major mode) recognition was only reduced by the left-resections. |
| 38 | *Ladinig & Schellenberg (2012)* | Liking unfamiliar music: effects of felt emotion and individual differences | High scores on Agreeableness and Neuroticism dimensions are associated with stronger sad feelings in response to music. Sad music (in minor mode) tends to be more appreciated among listeners who scored high Introversion or low Extroversion levels, or high Openness to experience. |
| 39 | *Leaver & Halpern (2004)* | Effects of training and melodic features on mode perception | Non-musicians found mode discrimination to be harder than discrimination of other melodic features. Trained non-musicians and musicians showed a better performance but not to ceiling levels. |
| 40 | *Lee et al. (2011)* | Investigation of melodic contour processing in the brain using multivariate pattern-based fMRI | In a fMRI study, ascending and descending melodic sequences in major mode are perceived as happier than minor melodies irrespective of the contour. Searchlight analysis with respect to mode (major vs. minor) did not find any significant voxels. |
| 41 | *Lin et al. (2014)* | Revealing spatio-spectral electroencephalographic dynamics of musical mode and tempo perception by independent component analysis | Empirical results showed that music in major mode increases delta band activity over the right sensorimotor cortex, decreases theta activity over the superior parietal cortex and moderately suppresses beta activity over the medial frontal cortex compared to minor-mode music, where fast tempo music only engages significant alpha suppression over the right sensorimotor cortex. |
| 42 | *Mao & Rau (2014)* | EEG-based measurement of emotion induced by mode, rhythm, and MV of Chinese pop music | Among Chinese listeners, major mode was significantly more positive, and of higher arousal level than that induced by minor mode. Moreover, major mode is more likely to activate people and evoke feelings of vitality, happiness, and excitement |
| 43 | *Maslennikova et al. (2013)* | Evoked changes in EEG band power on perception of consonant and dissonant chords | Subjective evaluations of major and minor chords elicited no significant differences in terms of pleasantness/unpleasantness. |
| 44 | *McConnel & Shore (2011)* | Upbeat and happy: Arousal as an important factor in studying attention | Participants were in a more pleasant mood after listening to music written in major mode compared to music written in minor mode. Furthermore, musical mode influenced affective valence and arousal. |
| 45 | *Mednicoff et al. (2018)* | Many listeners cannot discriminate major vs minor tone-scrambles regardless of presentation rate | Listeners' performance in a major/minor classification task is not depending on the speed of presentation. |
| 46 | *Mizuno & Sugishita (2007)* | Neural correlates underlying perception of tonality-related emotional contents | No brain regions were found significantly related to listening to sequences in a major mode than those in a minor mode or vice versa. The data also suggest that the bilateral inferior gyri and thalamus are involved in sensing the mode. |
| 47 | *Nemoto et al. (2010)* | fMRI measurement of brain activities to major and minor chords and cadence sequences | Superior temporal gyrus and posterior and anterior cingulate are shown to be activated more strongly by minor chords than by major chords. In general, minor chords activated larger areas of brain regions than major chords. |
| 48 | *Nieminen et al. (2012)* | The development of the aesthetic Experience of music: Preference, emotions, and beauty | Children preferred the piece in major mode to the one in minor. But only 8-9-year-olds found the minor pieces sadder than the major pieces, and the major pieces more beautiful than the piece in minor. |
| 49 | *Pallesen et al. (2003)* | Emotional connotations of major and minor musical chords in musically untrained listeners | Untrained listeners are able to recognize the happy/sad emotional connotations of major/minor mode. |

| | | | |
|---|---|---|---|
| *50* | *Pallesen et al. (2005)* | Emotion processing of major, minor, and dissonant chords: a functional magnetic resonance imaging study | Minor and dissonant chords, compared with major chords, elicited enhanced responses in several brain areas, including the amygdala, retrosplenial cortex, brain stem, and cerebellum. Findings did not show any consistent difference between musicians and non-musicians when they listened to major, minor and dissonant isolated chords. |
| *51* | *Pallesen et al. (2015)* | Experience drives synchronization: the phase and amplitude dynamics of neural oscillations to musical chords are differentially modulated by musical expertise | Musicians reported a stronger gamma-band activity in response to non-prototypical chords in musicians, while beta-band activity was stronger in non-musicians when they listened to minor chords. |
| *52* | *Pilcher et al. (2014)* | Da capo: A musical technique to evoke narrative recall | In a memory evocation task, the major key evoked experiences of celebrations and 'triumphs,' and pieces in minor keys evoked experiences of exams, sadness, and disappointment. |
| *53* | *Proverbio et al. (2020)* | Multimodal recognition of emotions in music and facial expressions | Negative musical fragments (in minor mode) elicited greater auditory N400 than positive musical fragments (major mode), especially over the right hemisphere. |
| *54* | *Putkinen et al. (2014)* | Enhanced development of auditory change detection in musically trained school-aged children: A longitudinal event-related potential study | School-aged children showed MMNs to minor chords among major chords, with musical training and older age enhancing this effect. |
| *55* | *Schellenberg et al. (2008)* | Liking for happy- and sad- sounding music: Effects of exposure | After a long and complex task, the participants did not show the typical preference for happy over sad music. |
| *56* | *Smit et al. (2022)* | Emotional responses in Papua New Guinea show negligible evidence for a universal effect of major versus minor music | In a cross-cultural study, the typical major/happy and minor/sad association is strongly associated with exposure to Western or Western-like music. |
| *57* | *Steinbeis & Koelsch (2011)* | Affective priming effects of musical sounds on the processing of word meaning | Both musically trained and untrained participants evaluated emotional words more quickly when they were congruent to the affective connotation of a preceding chord (major-happy/minor-sad) as compared to words incongruent to the preceding chord. Moreover, a larger N400 for incongruous target words compared to congruous target words for both musically trained and musically untrained participants was found. |
| *58* | *Suda et al. (2008)* | Emotional responses to music: towards scientific perspective on music therapy | Salivary cortisol levels showed a significant decrease among major mode listeners. Moreover, in response to major mode music, haemoglobin increased. |
| *59* | *Suzuki et al. (2008)* | Discrete cortical regions associated with the musical beauty of major and minor chords | Minor chords perceived as beautiful, strongly activated the right striatum, whereas the "beautiful" major chords elicited significant left middle temporal gyrus activity. |
| *60* | *Tabei (2015)* | Inferior Frontal Gyrus Activation Underlies the Perception of Emotions, While Precuneus Activation Underlies the Feeling of Emotions during Music Listening. | Bilateral inferior frontal gyri and the precuneus are important areas for the perception of the emotional content of music and for the emotional response evoked in the listener |
| *61* | *Thompson & Opfer (2014)* | Affective constraints on acquisition of musical concepts: Children's and adults' development of the major–minor distinction | Adults, 10-year-olds, and 5-year-olds rated affective quality of excerpts that differed by mode, tempo, pitch, and excerpts type and then categorized the excerpts by mode. The assessment of category |

| | | | |
|---|---|---|---|
| | | | judgments indicated performance was accurately predicted by subjects' association of affective valence with musical properties. |
| *62* | *Trochidis & Bigand (2013)* | Investigation of the effect of mode and tempo on emotional responses to music using EEG power asymmetry | Musical modes influence the valence of emotion with major mode being evaluated happier and more serene than minor and Locrian mode. Furthermore, in EEG frontal activity, major mode was associated with an increased alpha activation in the left hemisphere compared to minor modes, which increased activation in the right hemisphere. |
| *63* | *Tsang et al., (2001)* | Frontal EEG responses as a function of affective musical features. | Change from minor to major (happier) induced increased left frontal activation while change from major to minor (sadder) decreased left frontal activation |
| *64* | *van der Zwaag et al. (2011)* | Emotional and Psychophysiological responses to tempo, mode, and percussiveness | Minor mode music evoked higher arousal ratings than major mode music. |
| *65* | *Virtala et al. (2011)* | The preattentive processing of major vs. minor chords in the human brain: An event-related potential study | Musically untrained individuals can neurally categorise chords even preattentively. Modes (minor, major) and degree of consonance (dissonance, consonance) can be preattentively encoded. |
| *66* | *Virtala et al. (2012)* | Musical training facilitates the neural discrimination of major versus minor chords in 13-year-old children | Only musically trained children demonstrated MMNs to the major- minor contrast. |
| *67* | *Virtala et al. (2013)* | Newborn infants' auditory system is sensitive to Western music chord categories | Newborn infants are sensitive to Western music categorizations since they showed statistically significant MMRs responses to dissonant and minor chords in a context of major chords. |
| *68* | *Wang et al. (2018)* | Study on the impact of different music education on emotional regulation of adolescents based on EEG signals | Minor music can achieve better psychological regulation and relaxation compared to major music. |
| *69* | Zhou et al. (2019) | Impaired emotional processing of chords in congenital amusia: Electrophysiological and behavioral evidence | Participants without amusia (vs amusia) had faster reaction times and a larger N400 when judging the pleasantness of affectively congruous target images compared to affectively incongruous target images |

**Table S3.**

*Quality Assessment of All included Studies*[3]

| | Author (year) | 01 | 02 | 03 | 04 | 05 | 06 | 07 | 08 | 09 | 10 | 11 | 12 | 13 | 14 | Total Score |
|---|---|---|---|---|---|---|---|---|---|---|---|---|---|---|---|---|
| 1 | *Abe et al. (2017)* | 2 | 2 | 2 | 1 | NA | NA | NA | 2 | 2 | 2 | 2 | 2 | 2 | 2 | 0.95 |
| 2 | *Al'tmann et al. (2000)* | 2 | 2 | 1 | 1 | NA | NA | NA | 2 | 1 | 1 | 1 | 1 | 2 | 2 | 0.73 |
| 3 | *Bakker & Martin (2015)* | 2 | 2 | 2 | 2 | NA | NA | NA | 2 | 2 | 2 | 2 | 2 | 2 | 2 | 1.00 |
| 4 | *Bonetti & Costa (2016)* | 2 | 2 | 2 | 2 | NA | NA | NA | 2 | 2 | 2 | 2 | 2 | 2 | 2 | 1.00 |
| 5 | *Bonetti & Costa (2019)* | 2 | 2 | 2 | 2 | NA | NA | NA | 2 | 2 | 2 | 2 | 2 | 2 | 2 | 1.00 |
| 6 | *Bonetti et al. (2017)* | 2 | 2 | 2 | 2 | NA | NA | NA | 2 | 2 | 2 | 2 | 2 | 2 | 2 | 1.00 |
| 7 | *Brattico et al. (2009)* | 2 | 2 | 2 | 2 | NA | NA | NA | 2 | 1 | 2 | 2 | 2 | 2 | 2 | 0.95 |
| 8 | *Brattico et al. (2011)* | 2 | 2 | 2 | 2 | NA | NA | NA | 2 | 1 | 2 | 2 | 2 | 2 | 2 | 0.95 |
| 9 | *Brattico et al. (2016)* | 2 | 2 | 2 | 2 | NA | NA | NA | 2 | 2 | 2 | 2 | 2 | 2 | 2 | 1.00 |
| 10 | *Castro & Lima (2014)* | 2 | 2 | 2 | 2 | NA | NA | NA | 2 | 2 | 2 | 2 | 2 | 2 | 2 | 1.00 |
| 11 | *Chang et al. (2015)* | 2 | 2 | 1 | 1 | NA | NA | NA | 1 | 2 | 2 | 2 | 1 | 2 | 2 | 0.82 |
| 12 | Chubb et al. (2013) | 2 | 2 | 1 | 1 | NA | NA | NA | 2 | 2 | 2 | 2 | 1 | 2 | 2 | 0.86 |
| 13 | *Costa (2013)* | 2 | 2 | 2 | 2 | NA | NA | NA | 2 | 2 | 2 | 2 | 2 | 2 | 2 | 1.00 |
| 14 | *Costa et al. (2004)* | 2 | 2 | 2 | 2 | NA | NA | NA | 2 | 2 | 2 | 2 | 2 | 2 | 2 | 1.00 |
| 15 | *Dalla Bella et al. (2001)* | 2 | 2 | 2 | 2 | NA | NA | NA | 2 | 2 | 2 | 2 | 2 | 2 | 2 | 1.00 |
| 16 | *Dobrota & Ervegovac (2015)* | 2 | 2 | 2 | 1 | NA | NA | NA | 2 | 2 | 2 | 2 | 1 | 2 | 2 | 0.91 |
| 17 | *Eerola et al. (2013)* | 2 | 2 | 2 | 2 | NA | NA | NA | 2 | 2 | 2 | 2 | 2 | 2 | 2 | 1.00 |
| 18 | *Ellison et al. (2015)* | 2 | 2 | 2 | 1 | NA | NA | NA | 2 | 1 | 2 | 2 | 1 | 2 | 2 | 0.86 |
| 19 | *Fang et al. (2017)* | 2 | 2 | 2 | 2 | NA | NA | NA | 2 | 2 | 2 | 2 | 2 | 2 | 2 | 1.00 |
| 20 | *Franco et al. (2017)* | 2 | 2 | 2 | 2 | NA | NA | NA | 2 | 2 | 2 | 2 | 2 | 2 | 2 | 1.00 |
| 21 | *Gosselin et al. (2005)* | 2 | 2 | 2 | 2 | NA | NA | NA | 2 | 2 | 2 | 2 | 2 | 2 | 2 | 1.00 |
| 22 | *Green et al (2008)* | 2 | 2 | 2 | 1 | NA | NA | NA | 2 | 2 | 2 | 2 | 1 | 2 | 2 | 0.91 |
| 23 | *Gregory et al. (1996)* | 2 | 2 | 1 | 1 | NA | NA | NA | 2 | 2 | 2 | 1 | 2 | 2 | 2 | 0.86 |
| 24 | *Halpern (1984)* | 2 | 2 | 2 | 2 | NA | NA | NA | 2 | 2 | 2 | 2 | 1 | 2 | 2 | 0.95 |
| 25 | *Halpern et al. (1998)* | 2 | 2 | 2 | 2 | NA | NA | NA | 2 | 2 | 2 | 2 | 2 | 2 | 2 | 1.00 |
| 26 | *Halpern et al. (2008)* | 2 | 2 | 2 | 2 | NA | NA | NA | 2 | 1 | 2 | 2 | 2 | 2 | 2 | 0.95 |

| | | | | | | | | | | | | | | | | |
|---|---|---|---|---|---|---|---|---|---|---|---|---|---|---|---|---|
| 27 | *Heaton et al. (1999)* | 2 | 2 | 2 | 1 | NA | NA | NA | 2 | 1 | 2 | 2 | 1 | 2 | 2 | 0.86 |
| 28 | *Hevner (1935)* | 2 | 1 | 2 | 1 | NA | NA | NA | 2 | 2 | 1 | 1 | 1 | 1 | 2 | 0.73 |
| 29 | *Hoshino (1996)* | 2 | 2 | 1 | 2 | NA | NA | NA | 2 | 2 | 2 | 2 | 2 | 1 | 2 | 0.91 |
| 30 | *Hunter et al. (2008)* | 2 | 2 | 2 | 2 | NA | NA | NA | 2 | 2 | 2 | 2 | 2 | 1 | 2 | 0.95 |
| 31 | *Hunter et al. (2011)* | 2 | 2 | 1 | 1 | NA | NA | NA | 2 | 2 | 2 | 2 | 1 | 2 | 2 | 0.86 |
| 32 | *Jenni et al. (2017)* | 2 | 2 | 2 | 2 | NA | NA | NA | 2 | 2 | 2 | 2 | 2 | 2 | 2 | 1.00 |
| 33 | *Jeong et al. (2011)* | 2 | 2 | 2 | 2 | NA | NA | NA | 2 | 1 | 2 | 2 | 2 | 2 | 2 | 0.95 |
| 34 | *Jomori et al. (2013)* | 2 | 1 | 2 | 2 | NA | NA | NA | 1 | 1 | 2 | 2 | 1 | 2 | 2 | 0.82 |
| 35 | *Kastner & Crowder (1990)* | 2 | 2 | 2 | 1 | NA | NA | NA | 2 | 1 | 2 | 2 | 1 | 2 | 2 | 0.86 |
| 36 | *Khalfa et al. (2005)* | 2 | 2 | 2 | 1 | NA | NA | NA | 2 | 2 | 2 | 2 | 1 | 2 | 2 | 0.91 |
| 37 | *Khalfa et al. (2008)* | 2 | 2 | 2 | 2 | NA | NA | NA | 2 | 2 | 2 | 2 | 2 | 2 | 2 | 1.00 |
| 38 | *Ladinig & Schellenberg (2012)* | 2 | 2 | 2 | 2 | NA | NA | NA | 2 | 2 | 2 | 2 | 2 | 2 | 2 | 1.00 |
| 39 | *Leaver & Halpern (2004)* | 2 | 2 | 2 | 2 | NA | NA | NA | 2 | 1 | 2 | 2 | 2 | 2 | 2 | 0.95 |
| 40 | *Lee et al. (2011)* | 2 | 2 | 2 | 1 | NA | NA | NA | 2 | 1 | 2 | 2 | 1 | 2 | 2 | 0.86 |
| 41 | *Lin et al. (2014)* | 2 | 2 | 2 | 2 | NA | NA | NA | 2 | 2 | 2 | 2 | 1 | 2 | 2 | 0.95 |
| 42 | *Mao & Rau (2014)* | 2 | 2 | 2 | 1 | NA | NA | NA | 2 | 1 | 2 | 1 | 1 | 2 | 2 | 0.82 |
| 43 | *Maslennikova et al. (2013)* | 2 | 2 | 2 | 1 | NA | NA | NA | 2 | 1 | 2 | 2 | 1 | 1 | 2 | 0.82 |
| 44 | *McConnel & Shore (2011)* | 2 | 2 | 2 | 1 | NA | NA | NA | 2 | 2 | 2 | 2 | 1 | 2 | 2 | 0.91 |
| 45 | *Mednicoff et al. (2018)* | 2 | 2 | 2 | 2 | NA | NA | NA | 2 | 2 | 2 | 2 | 2 | 2 | 2 | 1.00 |
| 46 | *Mizuno & Sugishita (2007)* | 2 | 2 | 1 | 1 | NA | NA | NA | 1 | 2 | 1 | 1 | 1 | 1 | 1 | 0.64 |
| 47 | *Nemoto et al. (2010)* | 0 | 1 | 1 | 0 | NA | NA | NA | 1 | 1 | 1 | 0 | 0 | 1 | 1 | 0.32 |
| 48 | *Nieminen et al. (2012)* | 2 | 2 | 2 | 2 | NA | NA | NA | 2 | 2 | 2 | 2 | 2 | 2 | 2 | 1.00 |
| 49 | *Pallesen et al. (2003)* | 2 | 2 | 1 | 1 | NA | NA | NA | 2 | 1 | 2 | 1 | 1 | 1 | 2 | 0.73 |
| 50 | *Pallesen et al. (2005)* | 2 | 2 | 2 | 1 | NA | NA | NA | 2 | 2 | 1 | 1 | 1 | 1 | 2 | 0.77 |
| 51 | *Pallesen et al. (2015)* | 2 | 2 | 2 | 2 | NA | NA | NA | 2 | 2 | 2 | 2 | 2 | 2 | 2 | 1.00 |
| 52 | *Pilcher et al. (2014)* | 2 | 2 | 2 | 1 | NA | NA | NA | 2 | 1 | 2 | 1 | 1 | 2 | 2 | 0.82 |
| 53 | *Proverbio et al. (2020)* | 2 | 2 | 2 | 1 | NA | NA | NA | 2 | 1 | 2 | 2 | 1 | 2 | 2 | 0.86 |
| 54 | *Putkinen et al. (2014)* | 2 | 2 | 2 | 1 | NA | NA | NA | 2 | 2 | 2 | 2 | 2 | 2 | 2 | 0.95 |
| 55 | *Schellenberg et al. (2008)* | 2 | 2 | 2 | 1 | NA | NA | NA | 2 | 2 | 2 | 2 | 2 | 2 | 2 | 0.95 |
| 56 | *Smit et al. (2022)* | 2 | 2 | 2 | 1 | NA | NA | NA | 2 | 2 | 2 | 2 | 1 | 2 | 2 | 0.91 |
| 57 | *Steinbeis & Koelsch (2011)* | 2 | 2 | 2 | 2 | NA | NA | NA | 2 | 2 | 2 | 2 | 1 | 2 | 2 | 0.95 |

| | | | | | | | | | | | | | | | | |
|---|---|---|---|---|---|---|---|---|---|---|---|---|---|---|---|---|
| *58* | *Suda et al. (2008)* | 2 | 2 | 2 | 1 | NA | NA | NA | 2 | 1 | 2 | 2 | 1 | 2 | 2 | 0.86 |
| *59* | *Suzuki et al. (2008)* | 2 | 2 | 2 | 2 | NA | NA | NA | 2 | 2 | 2 | 2 | 1 | 1 | 2 | 0.91 |
| *60* | *Tabei (2015)* | 2 | 2 | 2 | 1 | NA | NA | NA | 2 | 2 | 2 | 2 | 2 | 2 | 2 | 0.95 |
| *61* | *Thompson & Opfer (2014)* | 2 | 2 | 1 | 1 | NA | NA | NA | 2 | 1 | 2 | 2 | 1 | 2 | 2 | 0.82 |
| *62* | *Trochidis & Bigand (2013)* | 2 | 2 | 2 | 1 | NA | NA | NA | 2 | 1 | 2 | 1 | 1 | 2 | 2 | 0.82 |
| *63* | *Tsang et al., (2001)* | 2 | 2 | 0 | 1 | NA | NA | NA | 2 | 2 | 1 | 1 | 1 | 2 | 2 | 0.73 |
| *64* | *van der Zwaag et al. (2011)* | 2 | 2 | 2 | 1 | NA | NA | NA | 2 | 1 | 2 | 2 | 1 | 2 | 2 | 0.86 |
| *65* | *Virtala et al. (2011)* | 2 | 2 | 2 | 1 | NA | NA | NA | 2 | 2 | 2 | 2 | 2 | 2 | 2 | 0.95 |
| *66* | *Virtala et al. (2012)* | 2 | 2 | 2 | 1 | NA | NA | NA | 2 | 2 | 2 | 2 | 2 | 2 | 2 | 0.95 |
| *67* | *Virtala et al. (2013)* | 2 | 2 | 2 | 1 | NA | NA | NA | 2 | 1 | 2 | 1 | 1 | 2 | 2 | 0.82 |
| *68* | *Wang et al. (2018)* | 2 | 1 | 1 | 1 | NA | NA | NA | 1 | 2 | 1 | 1 | 0 | 2 | 2 | 0.64 |
| *69* | Zhou et al. (2019) | 2 | 2 | 2 | 2 | NA | NA | NA | 1 | 1 | 2 | 2 | 2 | 2 | 2 | 0.91 |

A summary score was calculated for each paper by summing the total score obtained across relevant items and dividing by the total possible score (i.e.: 28 – (number of "n/a" x 2)). A score higher than 0.75 suggests high quality, between 0.55 and 0.75 suggests moderate quality, and lower than 0.55 suggests low quality in the study. Since no qualitative studies were included in this review, only the checklist for assessing the quality of quantitative studies was used. The review only included observational studies, so items 5, 6, and 7 were scored as not applicable (NA). The total scores for each study ranged from 0.32 to 1.0 with a mean of 0.9 ± 0.12.

| Checklist for assessing the quality of quantitative studies[3] | |
|---|---|
| 1 | Question / objective sufficiently described? |
| 2 | Study design evident and appropriate? |
| 3 | Method of subject/comparison group selection *or* source of information/input variables described and appropriate? |
| 4 | Subject (and comparison group, if applicable) characteristics sufficiently described? |
| 5 | If interventional and random allocation was possible, was it described? |
| 6 | If interventional and blinding of investigators was possible, was it reported? |
| 7 | If interventional and blinding of subjects was possible, was it reported? |
| 8 | Outcome and (if applicable) exposure measure(s) well defined and robust to measurement/misclassification bias? Means of assessment reported? |
| 9 | Sample size appropriate? |
| 10 | Analytic methods described/justified and appropriate? |
| 11 | Some estimate of variance is reported for the main results? |
| 12 | Controlled for confounding? |
| 13 | Results reported in sufficient detail? |
| 14 | Conclusions supported by the results? |

#### Table S4

*fMRI Acquisition and Analysis*

| | | | | | | | Structural | | | | Functional | | | | | | | |
|---|---|---|---|---|---|---|---|---|---|---|---|---|---|---|---|---|---|---|
| | **Author** | **Year** | **Teslas** | **MRI-system** | **MRI-model** | **Head-coil** | **T1 sequence** | **TR (ms)** | **TE (ms)** | **Voxel size (mm)** | **T2*sequence** | **TR (ms)** | **TE (ms)** | **Thickness (mm)** | **Acquisition Type** | **Analysis method** | **Analysis software** | **Foci** |
| 1 | Brattico | 2011 | 3 | General Electric | Signa | - | - | - | - | - | EPI | 3 | 32 | 4 | Interleaved | ANOVA | SPM8 | Y |
| 2 | Brattico | 2016 | 3 | General Electric | Signa | - | T1w | - | - | - | EPI | 3 | 32 | 4 | Interleaved | GLM | SPM8 | Y |
| 3 | Green | 2008 | 1.5 | General Electric | Signa | - | T1w | - | - | - | EPI | 2.7 | 40 | 5 | - | GLM | SPM5 | Y |
| 4 | Jeong | 2011 | 3 | General Electric | Signa | 8-channel | SPGR | 9.12 | 3.66 | 1x1x1 | EPI | 3 | 35 | 4 | - | GLM | SPM8 | Y |
| 5 | Khalfa | 2005 | 3 | Bruker | Medspec | - | T1w | - | - | 1x0.75x1.22 | STS | - | - | 4 | Interleaved | GLM | SPM99 | Y |
| 6 | Lee | 2011 | 3 | Philips | Intera Achieva | - | MPRAGE | - | - | 1z1z1 | EPI | 2 | 35 | 3 | - | MVPA | SPM5 | N |
| 7 | Mizuno | 2007 | 1.5 | General Electric | Signa | - | T1w | - | - | - | EPI | 2 | 50 | 6.5 | - | GLM | SPM99 | Y |
| 8 | Nemoto | 2010 | 1.5 | Hitachi | Stratis-2 | - | - | - | - | - | - | - | - | 5 | - | - | SPM8 | N |
| 9 | Pallesen | 2003 | - | - | - | - | - | - | - | - | EPI | - | - | - | - | ANOVA | - | N |
| 10 | Pallesen | 2005 | 1.5 | Siemens | Sonata | - | - | - | - | - | - | - | - | - | - | - | FSL | N |
| 11 | Suzuki | 2008 | PET | - | - | - | - | - | - | - | - | - | - | - | - | T-stats | SPM2 | Y |
| 12 | Tabei | 2015 | 1.5 | Siemens | Symphony | - | T1w | 2.2 | 3.93 | 1x1x1 | EPI | 4 | 50 | 3 | - | GLM | SPM5 | N |

**Table S5**

*Functional Characterization of ROIs as Included on BrainMap Platform Resulted from ALE Meta-analyses*

| | |
|---|---|
| 1. Major - Minor | |
| 1.1. STG-L BA22a | |
| Action | Execution, speech, imagination, observation |
| Cognition | Attention, language, phonology, semantics, speech, explicit memory, working memory, music, reasoning, somatic |
| Emotion | Negative, anger, fear, sadness, positive emotion, happiness |
| Interoception | Sexuality |
| Perception | Audition, pain |
| Paradigms | Cued explicit recall, encoding, face discrimination, film viewing, finger tapping/button pressing, flexion/extension, go/no-go, hand-eye coordination, lexical decision, music comprehension, music production, oddball discrimination, pain discrimination, passive listening, passive viewing, phonological discrimination, pitch discrimination, reading, reasoning/problem solving, recitation/repetition, semantic discrimination, sexual arousal, tone discrimination, visuospatial attention, word generation |
| 1.2. MedFG BA6 | |
| Action | Execution, speech, imagination, inhibition |
| Cognition | Attention, orthography, semantics, speech, working memory, music, reasoning, social cognition, somatic |
| Emotion | Embarrassment, fear, punishment, reward/gain |
| Interoception | Gastrointestinal/genitourinary, sexuality |
| Perception | Audition, olfaction, somesthesis, pain, vision, motion, shape |
| Paradigms | Affective pictures, chewing/swallowing, classical conditioning, competition/cooperation, counting/calculation, cued explicit recall, deception, delayed match to sample, emotion induction, film viewing, finger tapping/button pressing, flanker, flashing checkerboard, flexion/extension, gambling, go/no-go, grasping, imagined movement, isometric force lip pursing/tongue movement, micturition, music comprehension, music production, n-back, naming, oddball discrimination, orthographic discrimination, pain discrimination, passive listening, phonological discrimination, pitch discrimination, pursuit rotor/manual tracking, reading, recitation/repetition, reward, saccades, semantic discrimination, sequence recall/learning, sexual arousal, tone discrimination, transcranial magnetic stimulation, video games, visual object identification, visual pursuit/tracking, visuospatial attention, word generation |
| 1.3. TTG-R BA41 | |
| Action | Execution |
| Cognition | Attention, phonology, semantics, speech, syntax, working memory, music, reasoning, temporal |
| Emotion | Negative, positive, reward/gain |
| Interoception | - |
| Perception | Audition, somesthesis, pain, vision, motion, shape |
| Paradigms | Classical conditioning, driving, figurative language, finger tapping/button pressing, gambling, go/no-go, music comprehension, music production, orthographic discrimination, pain discrimination, passive listening, phonological discrimination, passive listening, pitch discrimination, reading, reasoning/problem solving, recitation/repetition, reward, semantic discrimination, syntactic discrimination, tactile discrimination, tone discrimination, transcranial magnetic stimulation, visual object identification, word generation |
| 1.4. CG-R BA31 | |
| Action | Execution, preparation |
| Cognition | Attention, semantics, speech, explicit memory, working memory, music, reasoning |
| Emotion | Anxiety, disgust, reward/gain |
| Interoception | Sexuality |
| Perception | Audition, shape |
| Paradigms | Affective pictures, cued explicit recall, deception, delay discounting, delayed match to sample, encoding, episodic recall, face discrimination, film viewing, finger tapping/button pressing, fixation, go/no-go, imagined objects, music comprehension, music production, reward, sexual arousal, Stroop, word generation |
| 1.5. CAU-R | |
| Action | Execution, imagination, inhibition, observation, preparation |
| Cognition | Attention, phonology, semantics, syntax, explicit memory, working memory, music, reasoning, social cognition |
| Emotion | Negative, happiness, reward/gain |
| Interoception | Sexuality, thermoregulation |
| Perception | Audition, somesthesis, pain, vision, colour, shape |
| Paradigms | Counting/calculation, cued explicit recall, deception, delay discounting, encoding, episodic recall, face discrimination, film viewing, finger tapping/button pressing, flexion/extension, gambling, go/no-go, imagined movement, isometric |

| | |
|---|---|
| | force, music comprehension, n-back, oddball discrimination, pain discrimination, passive listening, passive viewing, reward, sexual arousal, tactile discrimination, task switching, tone discrimination, visual object identification, visuospatial attention |

#### Table S6

*Meta-analytic Connectivity Modelling of ROIs Resulted from ALE Meta-analyses.*

| Cluster Number | Volume (mm3) | MNI coordinates x | y | z | ALE | P | Z | Label (Side Region BA) |
|---|---|---|---|---|---|---|---|---|
| 1. Major – Minor | | | | | | | | |
| 1.1. STG-L BA22a: 946 foci, 53 experiments, 890 subjects (x=-52, y=-14, z=0) | | | | | | | | |
| 1 | 20272 | 60 | -14 | 0 | 8E-02 | 4E-22 | 9.6 | R Superior Temporal Gyrus BA22 |
| 2 | 20040 | -52 | -14 | 0 | 2E-01 | 0E+00 | 22.0 | L Superior Temporal Gyrus BA22 |
| 3 | 1464 | -12 | -16 | 4 | 3E-02 | 9E-08 | 5.2 | L Thalamus |
| 4 | 1136 | 6 | 14 | 38 | 3E-02 | 2E-07 | 5.1 | R Cingulate Gyrus BA32 |
| 1.2. MedFG-R BA6: 1520 foci, 67 experiments, 978 subjects (x=6, y=-4, z=60) | | | | | | | | |
| 1 | 45784 | 6 | -2 | 60 | 3E-01 | 0E+00 | 23.1 | R Medial Frontal Gyrus BA6 |
| 2 | 33048 | 14 | -18 | 6 | 5E-02 | 4E-12 | 6.8 | R Thalamus |
| 3 | 1952 | -30 | -54 | -26 | 4E-02 | 2E-07 | 5.1 | L Cerebellum |
| 4 | 1496 | -32 | 24 | 0 | 4E-02 | 2E-07 | 5.1 | L Insula BA13 |
| 5 | 1488 | 2 | -60 | -16 | 3E-02 | 8E-07 | 4.8 | R Cerebellum |
| 1.3. TTG-R BA41: 739 foci, 40 experiments, 599 subjects (x=46, y=-24, z=10) | | | | | | | | |
| 1 | 17000 | 46 | -22 | 10 | 1E-01 | 0E+00 | 16.8 | R Transverse Temporal Gyrus BA41 |
| 2 | 11168 | -40 | -30 | 14 | 5E-02 | 2E-14 | 7.6 | L Superior Temporal Gyrus BA41 |
| 3 | 1648 | 56 | -6 | 34 | 3E-02 | 2E-08 | 5.5 | R Precentral Gyrus BA6 |
| 4 | 1536 | -12 | -18 | 0 | 3E-02 | 5E-08 | 5.3 | L Thalamus |
| 5 | 1280 | 16 | -22 | 0 | 3E-02 | 5E-07 | 4.9 | R Thalamus |
| 1.4. CG-R BA31: 303 foci, 14 experiments, 212 subjects (x=8, y=-28, z=40) | | | | | | | | |
| 1 | 3432 | 6 | -28 | 38 | 7E-02 | 4E-28 | 10.9 | R Cingulate Gyrus BA31 |
| 1.5. CAU-R: 680 foci, 30 experiments, 511 subjects (x=20, y=-2, z=22) | | | | | | | | |
| 1 | 6960 | 20 | -2 | 22 | 1E-01 | 0E+00 | 16.9 | R Caudate |
| 2 | 2296 | 2 | 10 | 54 | 3E-02 | 1E-07 | 5.2 | L Superior Frontal Gyrus BA6 |
| 3 | 1872 | -30 | 18 | 10 | 2E-02 | 7E-06 | 4.4 | L Claustrum |
| 2. Minor – Major | | | | | | | | |
| 2.1. STG-L BA22b: 763 foci, 48 experiments, 709 subjects (x=-60, y=-8, z=-6) | | | | | | | | |
| 1 | 17544 | -60 | -8 | -6 | 2E-01 | 0E+00 | 20.6 | L Superior Temporal Gyrus BA22 |
| 2 | 11904 | 64 | -10 | -2 | 5E-02 | 9E-16 | 8.0 | R Superior Temporal Gyrus BA22 |
| 3 | 1056 | -56 | 10 | 10 | 2E-02 | 2E-05 | 4.2 | L Precentral Gyrus BA44 |
| 2.2. STG-R BA22: 1045 foci, 53 experiments, 776 subjects (x=60, y=-8, z=-2) | | | | | | | | |
| 1 | 18784 | -56 | -14 | 0 | 8E-02 | 2E-25 | 10.4 | L Superior Temporal Gyrus BA22 |
| 2 | 16304 | 60 | -8 | -2 | 2E-01 | 0E+00 | 21.5 | R Superior Temporal Gyrus BA22 |
| 3 | 2096 | -2 | 0 | 64 | 3E-02 | 4E-06 | 4.5 | L Medial Frontal Gyrus BA6 |
| 4 | 1456 | -52 | 20 | 26 | 3E-02 | 9E-07 | 4.8 | L Middle Frontal Gyrus BA9 |
| 5 | 1184 | -36 | 22 | 0 | 3E-02 | 3E-07 | 5.0 | L Insula BA13 |

ALE, anatomic likelihood estimation; M, musicians; NM, non-musicians; GM, gray matter; WM, white matter; BA, Brodmann area; ROIs, regions-of-interest; P, p-value; Z, peak z-value; R, right; L, left. **ROIs**: IFG, inferior frontal gyrus; IPL, inferior parietal lobule; IC, internal capsule; PostCG, postcentral gyrus (primary somatosensory cortex or S1); PreCG, precentral gyrus (primary motor cortex or M1); STG, superior temporal gyrus (primary auditory cortex). Music-related ROIs were created in Mango (http://rii.uthscsa.edu/mango//userguide.html) with a 5mm-radius sphere. Last search in Sleuth, 09.01.2023 (http://www.brainmap.org/sleuth/); NA, not enough available observations.

| 2. Minor – Major | |
|---|---|
| 2.1. STG-L BA22b | |
| Action | Execution, speech, imagination, observation |
| Cognition | Attention, language, orthography, phonology, semantics, speech, syntax, explicit memory, working memory, music, reasoning, social cognition, somatic |
| Emotion | Fear, sadness, positive, happiness, reward/gain, valence |
| Interoception | Thermoregulation |
| Perception | Audition, pain, vision, motion, shape |
| Paradigms | Affective pictures, counting/calculation, divided auditory attention, emotion induction, face discrimination, figurative language, film viewing, finger tapping/button pressing, fixation, imagined objects, lexical decision, multi-tasking, music comprehension, n-back, naming, orthographic discrimination, pain discrimination, paired associate recall, passive listening, passive viewing, phonological discrimination, pitch discrimination, reading, reasoning/problem solving, reward, saccades, semantic discrimination, sequence recall, Stroop, syntactic discrimination, task switching, theory-of-mind, tone discrimination |
| 2.2. STG-R BA22 | |
| Action | Execution, speech, observation |
| Cognition | Attention, language, phonology, semantics, speech, syntax, working memory, music, reasoning |
| Emotion | Fear, happiness, reward/gain, valence |
| Interoception | Sexuality |
| Perception | Audition, gustation, somesthesis, vision |
| Paradigms | Affective pictures, counting/calculation, divided auditory attention, emotion induction, face discrimination, figurative language, film viewing, finger tapping/button pressing, fixation, imagined objects, lexical decision, multi-tasking, music comprehension, n-back, naming, orthographic discrimination, pain discrimination, paired associate recall, passive listening, passive viewing, phonological discrimination, pitch discrimination, reading, reasoning/problem solving, reward, saccades, semantic discrimination, sequence recall, Stroop, syntactic discrimination, task switching, theory-of-mind, tone discrimination |

ALE, anatomic likelihood estimation; M, musicians; NM, non-musicians; GM, grey matter; WM, white matter; BA, Brodmann area; ROIs, regions-of-interest; P, p-value; Z, peak z-value; R, right; L, left. **ROIs**: IFG, inferior frontal gyrus; IPL, inferior parietal lobule; IC, internal capsule; PostCG, postcentral gyrus (primary somatosensory cortex or S1); PreCG, precentral gyrus (primary motor cortex or M1); STG, superior temporal gyrus (primary auditory cortex). Music-related ROIs were created in Mango (http://rii.uthscsa.edu/mango//userguide.html) with a 5mm-radius sphere. Last search in Sleuth, 09.01.2023 (http://www.brainmap.org/sleuth/); NA, not enough available observations.

*Table S7*

*FSN Robustness Assessment of ROIs Resulted from ALE Meta-analyses*

| Cluster number | Volume (mm³) | MNI coordinates x | y | z | ALE | Label (Side, region) | FSN |
|---|---|---|---|---|---|---|---|
| 1. Major – Minor: 65 foci, 7 experiments, 119 subjects, minimum FSN = 2 | | | | | | | |
| 1 | 1272 | -52 | -14 | 0 | 2E-02 | L Superior Temporal Gyrus BA22 | 6 |
| 2 | 824 | 6 | -4 | 60 | 2E-02 | R Medial Frontal Gyrus BA6 | 4 |
| 3 | 808 | 46 | -24 | 10 | 2E-02 | R Transverse Temporal Gyrus BA41 | 2 |
| 4 | 768 | 8 | -28 | 40 | 2E-02 | R Cingulate Gyrus BA31 | 2 |
| 5 | 488 | 20 | -2 | 22 | 2E-02 | R Caudate | <2 |
| 2. Minor – Mayor: 33 foci, 8 experiments, 138 subjects, minimum FSN = 2 | | | | | | | |
| 1 | 1096 | -60 | -8 | -6 | 2E-02 | L Superior Temporal Gyrus BA22 | 4 |
| 2 | 1040 | 60 | -8 | -2 | 2E-02 | R Superior Temporal Gyrus BA22 | 4 |

FSN, Fail-Safe N analysis; NA, not enough available observations.
